# Crystal structure of the *Legionella pneumophila* effector SidL (Lpg0437) in complex with its metaeffector LegA11 (Lpg0436)

**DOI:** 10.1101/2025.10.23.683979

**Authors:** Dominik A. Machtens, Carissa A. Hutchison, Ashley M. Stein, Jan Eberhage, Jonas M. Willerding, Susanne Eschenburg, Stephanie R. Shames, Thomas F. Reubold

**Author notes:** Department of Gastroenterology, Hepatology, Infectious Diseases and Endocrinology, Hannover Medical School, Carl-Neuberg-Straße 1, 306025 Hannover, Germany (Dominik A. Machtens). Department of Nephrology and Hypertension, Hannover Medical School, Carl-Neuberg-Straße 1, 306025 Hannover, Germany (Jonas M. Willerding). Denotes equal contribution (Carissa A. Hutchison, Ashley M. Stein, Stephanie R. Shames, Thomas F. Reubold). **CONTACT** Stephanie R. Shames; Thomas F. Reubold.

## Abstract

*Legionella pneumophila* is an opportunistic human pathogen that causes atypical pneumonia called Legionnaires’ Disease. To replicate within host cells, *L. pneumophila* injects up to 330 effector proteins into the host cytosol via a Dot/Icm type IV secretion system. Several effectors, called metaeffectors, regulate the activity of other effectors within infected host cells through direct protein-protein interactions. LegA11 (AnkJ/Lpg0436) has been identified as a putative metaeffector of SidL (Ceg14/Lpg0437), one of eight *L. pneumophila* effectors that inhibit host mRNA translation. LegA11 binds and suppresses SidL enzymatic activity, but the molecular basis of this regulation and impact on mRNA translation are unknown. Here, we present the crystal structure of SidL in complex with LegA11 to a resolution of 2.4 Å, revealing a high-affinity 1:1 complex with an extensive interaction interface of ∼2300 Å². Using isothermal titration calorimetry, we determined a dissociation constant of 1.8 nM. *In vitro* translation assays demonstrate that SidL inhibits mRNA translation, and this activity is completely suppressed by LegA11. Mutagenesis of key interface residues in LegA11 disrupts complex formation and abolishes its metaeffector activity, confirming that LegA11 regulates SidL through direct protein-protein interaction. These findings establish LegA11 as a *bona fide* metaeffector that contributes to suppression of host mRNA translation by *L. pneumophila*.

## Introduction

*Legionella pneumophila* is an opportunistic pathogen that is ubiquitous in freshwater environments where it parasitizes free-living amoebae [1]. *L. pneumophila* can also replicate within alveolar macrophages, leading to a potentially life-threatening atypical pneumonia called Legionnaires’ Disease. To replicate within host cells, *L. pneumophila* remodels the phagosome into a replication-permissive endoplasmic reticulum-like compartment called the *Legionella*-containing vacuole (LCV). To achieve this, *L. pneumophila* injects up to 330 effector proteins into the cytosol of the host cell through a Dot/Icm type IV secretion system (T4SS) [2]. *L. pneumophila* effectors interfere with and subvert key host cellular processes including ubiquitin-dependent proteasomal degradation, cell death, phagocytosis, vesicular trafficking, autophagy, and mRNA translation [3].

Several cellular processes are targeted by multiple effectors, reflecting *L. pneumophila*’s co-evolution with phylogenetically diverse hosts [4]. So far, eight different effectors have been identified in *L. pneumophila* that inhibit host mRNA translation. Lgt1 (Lpg1368), Lgt2 (Lpg2862), and Lgt3 (Lpg1488) are glycosyl transferases that transfer a glucose moiety to a specific serine residue in the eukaryotic translation elongation factor 1A [5,6]. The serine/threonine kinase LegK4 (Lpg0208) blocks translation by phosphorylation of a threonine residue in Hsp70 proteins [7], whereas VipF (Lpg0103) is an acetyl transferase that targets eukaryotic initiation factor 3 complex [8]. SidI (Lpg2504) transfers a mannosyl group to several ribosomal proteins [9]. The mechanism of action of RavX (Lpg1489) is currently unknown. SidL (Ceg14/Lpg0437) blocks translation via an unknown mechanism but was recently shown to function as an ATPase that hydrolyzes ATP to AMP and pyrophosphate in the presence of host actin (He et al., 2025). The dedication of multiple effectors to subversion of host translation machinery underscores its importance for *L. pneumophila* virulence.

Work over the past 15 years has shown that a subset of *L. pneumophila* effectors, termed metaeffectors, do not exclusively target host proteins but instead serve to regulate the function of other effectors through a direct protein-protein interaction [10,11], distinguishing them from antagonistic effectors that exert opposing biochemical activities on the same host factor(s) [11]. Both SidI and SidL are regulated by metaeffectors, suggesting an important role for this regulatory mechanism in *L. pneumophila*’s subversion of host mRNA translation. SidI is regulated by MesI (<u>Me</u>taeffector of <u>S</u>idI), which forms a high-affinity 1:1 complex with SidI and fully suppresses SidI-mediated translation inhibition and toxicity [12,13]. The effector LegA11 (AnkJ/Lpg0437) binds SidL and alleviates the toxic effect of SidL in yeast cells [14,15], but the molecular and biophysical underpinnings of the SidL-LegA11 interaction and impact of LegA11 on SidL-mediated translation inhibition are unknown.

Here, we present the crystal structure of SidL in complex with its cognate metaeffector LegA11 to a resolution of 2.4 Å. Using an *in vitro* translation assay, we show that SidL inhibits mRNA translation by an order of magnitude and that this activity is suppressed in the presence of LegA11. Based on the crystal structure, we identify residues in LegA11 that upon mutation disrupt the binding interface and abolish the metaeffector activity of LegA11. Thus, LegA11 is a *bona fide* metaeffector that contributes to *L. pneumophila*’s subversion of host mRNA translation.

## Material and Methods

### Bacterial strains, cell culture, and growth conditions

*Escherichia coli* strains used for cloning (Top10; DH5α*pir*) and protein expression [BL21 (DE3)] were grown at 37 °C in lysogeny broth (LB) medium supplemented with antibiotics for plasmid selection (100 µg/mL ampicillin, 50 µg/mL kanamycin, 25 µg/mL chloramphenicol). *L. pneumophila* SRS43 (Shames et al., 2017) was cultured on ACES-buffered charcoal yeast extract (CYE) agar or grown in ACES-buffered yeast extract (AYE) freshly supplemented with L-cysteine and ferric pyrophosphate at 37 °C, as described [16]. When necessary, media were supplemented with 10 µg/mL chloramphenicol for plasmid selection. Plasmid-based gene expression in *L. pneumophila* was induced with 1 mM IPTG (GoldBio). *Acanthamoeba castellanii* strain Neff was acquired from the American Type Culture Collection (ATCC #30010) and maintained in PYG broth [7.5 g/L proteose peptone, 7.5 g/L yeast extract, 15 g/L D(+)-glucose, 0.4 mM CaCl2, 4 mM MgSO4, 3.4 mM Na citrate, 50 µM F3(NH4)2(SO4)2, 2.5 mM Na2HPO4, 2.5 mM KH2PO4, pH 6.5], as described [17].

### Molecular cloning and strain construction

Oligonucleotide primer sequences are listed in Suppl. Table 1. For expression of His6-Myc fusion proteins, the respective *L. pneumophila* genes were cloned into pT7HMT, a gift from Dr. Brian Geisbrecht [18]. Wild-type *sidL* and *legA11* were amplified from *L. pneumophila* SRS43 genomic DNA (gDNA) using SidLBamHI-F/SidLNotI-R and LegA11BamHI-F/LegA11NotI-R primer pairs. *legA11*K195A/F239R/G240R/F243R (*legA11*-4M) was amplified from a custom high-copy cloning vector purchased from Twist Biosciences (San Francisco, CA) using LegA11BamHI-F/LegA11NotI-R. BamHI/NotI-digested fragments were purified and ligated into BamHI/NotI-digested pT7HMT [18]. Sequence-confirmed constructs (Genewiz) were transformed into chemically competent *E. coli* Top10 and BL21 (DE3) for maintenance and protein expression, respectively. For genetic complementation of Δ*sidL* and Δ*legA11* mutations, *sidL* and *legA11* were amplified from *L. pneumophila* SRS43 gDNA using SidLJBBamHI-F/SidLSalI-R and LegA11JBBamHI-F/LegA11SalI-R primer pairs and ligated as BamHI/SalI fragments into BamHI/SalI-digested pSN85, a gift from Dr. Criag Roy [19]. Sequence confirmed pSN85::*legA11* and pSN85::*sidL* plasmids were transformed into electrocompetent *L. pneumophila* Δ*legA11* and Δ*sidL*, respectively, as described [12].

*L. pneumophila* SRS43 Δ*legA11* and Δ*sidL* were generated as described previously [20]. Deletion constructs were generated by cloning 5′ and 3′ flanking regions of each gene into pSR47S. To generate pSR47S::Δ*legA11,* 5′ and 3′ flanking regions were amplified using LegA11KO1-F/LegA11KO1-R and LegA11KO2-F/LegA11KO2-R primer pairs, digested with SacI/NotI and NotI/SalI, respectively, and ligated into SacI/SalI-digested pSR47S, a gift from Dr. Craig Roy. To generate pSR47S::Δ*sidL,* 5′ and 3′ flanking regions were amplified using SidLKO1-F/SidLKO1-R and SidLKO2-F/SidLKO2-R primer pairs, digested with SacI/NotI and NotI/SalI, respectively, and ligated into SacI/SalI-digested pSR47S. Ligation reactions were transformed into chemically competent *E. coli* DH5α*pir.* Deletion constructs were validated by colony PCR and Sanger Sequencing (Genewiz) and conjugated into *L. pneumophila* SRS43 for allelic exchange, as described previously [21]. Sucrose-resistant, kanamycin-sensitive colonies were screened by PCR and Sanger sequencing to verify gene deletion.

### Protein expression and purification

The coding sequences of *sidL* (*lpg0437*) (UniProt accession number Q5ZYD5) in full-length (SidL_1s_BamHI/SidL_666as_SalI) or in N- and C-terminally truncated form (SidL_23s_BamHI/SdiL_640as_SalI; SidL_76s_BamH/SidL_645as_SalI) and of *legA11* (*lpg0436*) in full-length (LegA11_1s_BamHI/LegA11_269as_Sal/) were amplified from *Legionella pneumophila* strain Philadelphia 1 (DSM-7513, DSMZ, Braunschweig, Germany) gDNA using primer pairs and cloned into either pGEX-4T1-TEV or pMEX-4T1-TEV [22]. Point mutations were introduced by overlap-PCR. The correctness of the resulting constructs was verified by sequencing (Seqlab, Göttingen, Germany). The constructs were expressed in *Escherichia coli* BL21 (DE3) cells grown to an OD of 1.0 in terrific broth. Expression was induced with 0.2 mM IPTG overnight at 293 K. The bacteria were harvested by centrifugation for 15 min at 5000 × g. Pellets were resuspended in 50 mM HEPES pH 7.5, 200 mM NaCl, 5 mM 2-mercaptoethanol, 1 mM PMSF, 0.5% (v/v) NP-40, and 0.3 mg/mL hen egg-white lysozyme and incubated on ice for 30 min. The cells were lysed by sonication and cellular debris was removed by centrifugation at 32 000 × g for 30 min. The supernatant was loaded onto either an amylose sepharose or a GST column, depending on the respective purification tag, equilibrated with a buffer containing 50 mM HEPES pH 7.5, 300 mM NaCl, and 5 mM 2-mercaptoethanol. After washing with 10 column volumes of buffer the bound fusion protein was digested on column with TEV protease overnight. The cleaved protein was eluted with column buffer, concentrated to a volume of 2 mL, and applied to a S200 16/600 size exclusion column (GE Healthcare) equilibrated with a buffer containing 20 mM HEPES pH 7.5, 150 mM NaCl, and 2 mM DTT. The peak fractions were pooled, concentrated and flash-frozen in liquid nitrogen. For preparation of seleno-methionine (SeMet) substituted SidL and LegA11, proteins were expressed in *E. coli* BL21(DE3) following a protocol using PSM-5052 auto-induction medium [23].

### Analytical size exclusion chromatography

Equimolar amounts of SidL and/or LegA11 in a total volume of 100 µL were applied to an S200 10/300 size exclusion column (GE Healthcare) equilibrated with buffer containing 20 mM HEPES pH 7.5, 150 mM NaCl, and 2 mM DTT at a flow rate of 0.5 ml/min. The concentrations used are given in the respective figure legends. The figures representing the chromatographic data were prepared using Origin (Version 2022b. OriginLab Corporation, Northampton, MA, USA).

### Crystallization

Crystallization screening using various commercially available screens was performed in the 96-well format by mixing 200 nL of protein solution containing 12 mg/mL SidL23-640, 5.2 mg/mL LegA11, and chymotrypsin, trypsin, or thermolysin in ratios between 1:500 to 1:2000 with 200 nL of reservoir solution in MRC plates using a Phoenix crystallization robot (Art Robbins Instruments, Sunnyvale, USA). The droplets were equilibrated against 50 µL reservoir solution at 291 K. Plate-shaped crystals grew in condition F1 (100 mM HEPES-NaOH pH 7.5, 200 mM L-proline, 100 g/L PEG3350) of the Classics 2 screen (Qiagen, Hilden, Germany) in the presence of trypsin (T5266, Sigma-Aldrich, Taufkirchen, Germany) at 17 µg/mL. SeMet-derivatized crystals used for cryogenic data collection were grown in droplets of 1 µL protein (12 mg/mL SidL23-640, 5.2 mg/mL LegA11, trypsin at 34 µg/mL) and 1 µL reservoir containing 100 mM HEPES-NaOH pH 7.5, 200 mM L-proline, and 200 g/L PEG3350 equilibrated against 200 µl reservoir solution. Crystal were cryoprotected by brief immersion in reservoir solution containing 140 g/L PEG3350 and 23% (v/v) ethylene glycol and flash-cooled in liquid nitrogen. Native crystals used for cryogenic data collection were grown in droplets of 1 µL protein (12 mg/mL SidL23-640, 5.2 mg/mL LegA11, trypsin at 340 µg/mL) and 1 µL reservoir containing 100 mM HEPES-NaOH pH 7.5, 200 mM L-proline, and 140 g/L PEG3350 equilibrated against 200 µl reservoir solution. Crystals were cryoprotected by serial immersion in reservoir solutions containing 140 g/L PEG3350 and 0, 2, 4, 7, 10, 13, 16, 19, or 23% (v/v) glycerol and flash-cooled in liquid nitrogen.

### Data collection and processing

A dataset from a single native crystal was collected to a resolution of 2.35 Å at beamline PROXIMA 2A at the synchrotron SOLEIL (Gif-sur-Yvette Cedex, France) using a wavelength of 0.9762 Å. Four datasets were collected from a crystal grown from SeMet-substituted protein at a wavelength of 0.9795 Å at beamline P13 at the PETRA III storage ring (Hamburg, Germany) and merged to yield a complete highly redundant dataset to 3.3 Å. Data were processed with XDS and scaled with XSCALE [24,25]. For both crystals space group P212121 could be assigned with unit cell dimensions of a = 68.5 Å, b = 93.4 Å, and c = 133.7 Å for the native crystal and of a = 65.7 Å, b = 97.4 Å, and c = 130.7 Å for the SeMet-crystal, respectively.

### Structure solution and refinement

The structure was solved by the single anomalous dispersion (SAD) method employing the CRANK2 pipeline [26]. The initial electron density map was of sufficient quality for automatic building of 614 residues. The structure was iteratively completed and refined against the native data using Phenix.refine [27] followed by visual inspection and adjustment of the model in COOT [28] after each refinement cycle. Images showing structural data were prepared using Pymol (The PyMOL Molecular Graphics System, Version 1.8.6.2, Schrödinger, LLC). The data statistics are summarized in Table 1. The atomic coordinates and structure factors of SidL/LegA11 have been deposited in the Protein Data Bank (PDB entry 9SXC).

**Table 1.**
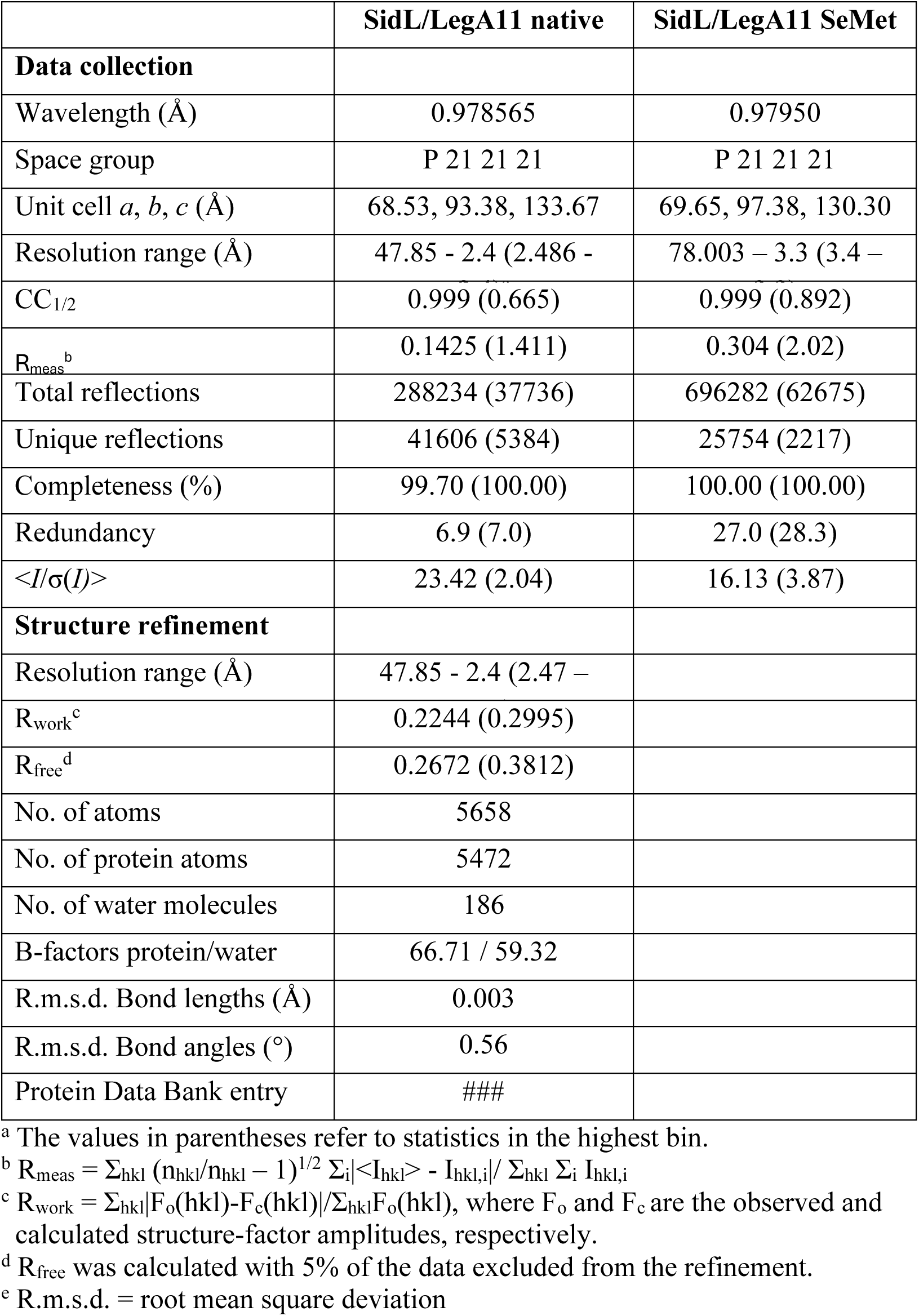
Data collection and refinement statistics.

### Isothermal titration calorimetry (ITC)

Binding between SidL (residues 76-645) and LegA11 (residues 1-269) was measured using a NanoITC 5302 microcalorimeter (Waters/TA Instruments) at 25 °C. Proteins were prepared in 20 mM HEPES (pH 7.5), 150 mM NaCl, degassed for 15 min at 30 mbar, and centrifuged at 20 000 × g for 5 min before titration. The sample cell (971 µL) contained SidL at 0.6 to 1.5 µM, with concentrations determined photometrically at 280 nm for each experiment. The syringe (100 µL) was loaded with 10 to 30 µM LegA11, also quantified by UV absorbance. Each experiment consisted of 20 injections of 4.91 µL ligand, delivered at 200 rpm stirring speed with 400 s spacing. Buffer-into-protein control titrations were performed, and the resulting heats were subtracted from binding data. Thermograms were integrated using the NanoITC software, and binding isotherms were fitted in Origin (OriginLab) with an independent binding model. Parameters obtained directly from the fit included the stoichiometry (*n*), dissociation constant (*K*d), and enthalpy change (Δ*H*); Gibbs free energy (Δ*G*), entropy (Δ*S*), and −*T*Δ*S* were calculated subsequently. Errors were estimated by error propagation, with asymmetric standard deviations reported for *K*d. Data are representative of four independent experiments.

### *In vitro* translation assay

Recombinant His6-Myc-SidL and -LegA11 were expressed in *E. coli* BL21 (DE3) as described [12]. Bacteria were pelleted by centrifugation at 2100 × g for 10 min at 4 °C and washed in 5 mL of ice-cold PBS. Bacterial pellets were resuspended in ice-cold lysis buffer [His Binding buffer (Zymo His-Spin Protein Miniprep Kit),10 µg/mL lysozyme (Sigma, Burlington, MA), and 10 mM 2-mercaptoethanol] and incubated on ice for 2-4 h followed by sonication using Kontes Micro Ultrasonic Cell Disrupter (40 amplitude, 30 s, five times). Lysates were clarified via centrifugation at 9600 × g for 15 min at 4 °C. Supernatants were transferred to clean microcentrifuge tubes. His-tagged proteins were then purified utilizing Zymo His-Spin Miniprep kit via manufacturer’s instructions (Zymo Research). Purity was validated by SDS-PAGE and Coomassie Blue staining. Purified recombinant proteins were dialyzed overnight in PBS pH 7.4 at 4 °C and quantified using Coomassie Plus Bradford Assay (BioRad) reagent on a BioTek Epoch2 microplate reader (Agilent).

Translation of *Firefly* Luciferase mRNA in Rabbit Reticulocyte Lysates was quantified using the Flexi Rabbit Reticulocyte Kit (Promega, Madison, WI) as described previously [12]. Purified recombinant His6-Myc-SidL (50 ng) and/or His6-Myc-LegA11 (23.3 ng) were added to lysates in equimolar amounts (65 nM). Lysates were supplemented with 20 µM ATP/Mg^2+^, as indicated. Reactions were run in triplicate for 90 min at 30 °C followed by addition of Luciferase assay reagent (Promega) and luminescence was quantified (arbitrary units) using a BioTek Synergy HTX multimode plate reader (Agilent).

### L. pneumophila *growth curve*s

*Legionella pneumophila* replication within *A. castellanii* was quantified as previously described [20]. One day prior to infection, cells were split 1:5 into 75 cm2 tissue culture flasks and incubated in PYG media at 25 °C. Cells were seeded in 24-well tissue culture-treated dishes (2.5 x 10^5^ cells/well) in 1 mL PYG and incubated at 37 °C for 2h prior to infection. Cells were washed in pre-warmed Ac buffer (0.4 mM CaCl2, 4 mM MgSO4, 3.4 mM Na citrate, 0.05 mM F3(NH4)2(SO4)2, 2.5 mM Na2HPO4, 2.5 mM KH2PO4, pH 6.5) and infected with *L. pneumophila* in 1 mL Ac buffer in triplicate wells at a multiplicity of infection (MOI) of 0.1. Cells were incubated for 45 min to allow bacterial uptake. Monolayers were washed three times to remove extracellular bacteria and either lysed or incubated in 1 mL Ac buffer for up to 72h at 37 °C. Cells were lysed in sterile deionized water and lysates were plated on CYE agar for enumeration of colony forming units (CFU) at 45 min, 24 h, 48 h, and 72 h post-infection.

To quantify *L. pneumophila* growth *in vitro*, 2-day heavy patches grown on CYE agar were resuspended in supplemented AYE broth and subcultured to an OD600 of 0.2. Cultures were dispensed into sterile 96-well round bottom plates (n=6/strain) and incubated with continuous orbital shaking in a BioTek Epoch2 microplate reader. Data were collected every 2h for up to 46h.

### Statistical analyses

Data are shown as mean ± standard deviation (s.d.) or standard error (s.e.m), as indicated. Statistical analyses were performed with GraphPad Prism 10 using either Welch’s *t*-test, one-way ANOVA, or two-way ANOVA, as described, with statistical significance defined as *P*<0.05. Unless otherwise indicated, statistical analyses were performed on biological triplicate samples (N=3) and representative of three independent experiments.

## Results

### SidL directly interacts with LegA11 forming a stable 1:1 complex

LegA11 was initially identified as a putative metaeffector based on its ability to suppress the toxic effect of SidL in a heterologous yeast expression system and bind to SidL in yeast two-hybrid and LUMIER assays [14]. To verify the direct interaction of SidL and LegA11, we subjected purified recombinant full-length SidL1-666, LegA11, or equimolar amounts of SidL1-666 and LegA11 to analytical size exclusion chromatography (SEC). In the chromatogram corresponding to the mixture, the peaks of the single proteins have merged into a single peak shifted to a lower elution volume (Fig. 1). This indicates that SidL and LegA11 form a stable complex with 1:1 stoichiometry.

**Figure 1.**
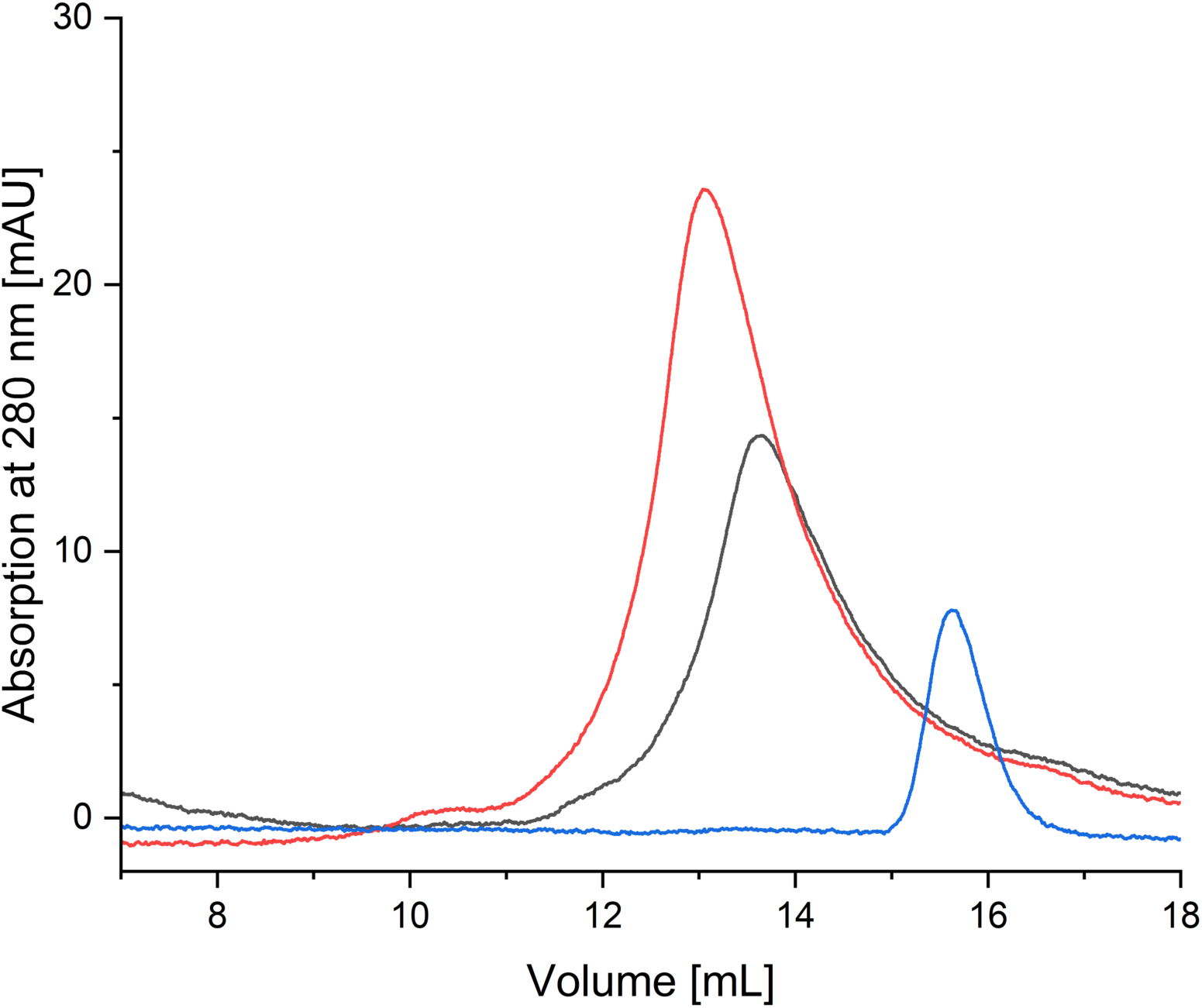
Interaction of SidL and LegA11 observed in analytical size exclusion chromatography. 300 µg SidL1-666 and/or 125 µg LegA11 in 100 µL buffer were applied to an S200 10/300 increase size exclusion column. The absorption curve of SidL1-666 is shown in black, the absorption curve of LegA11 is shown in blue, the absorption curve of the mixed proteins is shown in red.

We employed X-ray crystallography to visualize the SidL/LegA11 complex. Full-length SidL could not be purified in amounts large enough for an extended crystallization campaign; thus, we used the truncated variant SidL23-640 for overexpression as it could be produced in higher yield and displayed higher stability than the full-length protein. We set up crystallization trials using various commercial screens with an equimolar mixture of SidL and LegA11 but did not obtain any crystals. We then used *in situ* proteolysis [29] by adding various proteases to the crystallization setups (see *Materials and Methods*). We obtained crystals in several conditions, the best of which diffracted X-rays to a resolution limit of 2.4 Å after optimization. We solved the structure by SAD using data from a crystal composed of selenomethionine-substituted protein. Crystallographic details are given in Table 1.

The crystal structure contains residues 76-313, 334-413, and 525-628 of SidL and residues 7-192, and 202-261 of LegA11, as well as 186 water molecules. SidL is divided into two subdomains comprising residues 76-222 (SidLN) and residues 228-628 (SidLC), respectively (Fig. 2). SidLN is connected to SidLC via a straight elongated stretch of five residues (223-227). The two subdomains are separated by a C-terminal portion of LegA11 and do not interact with each other directly. Presumably, in the absence of LegA11 the domains display a high degree of conformational flexibility.

**Figure 2.**
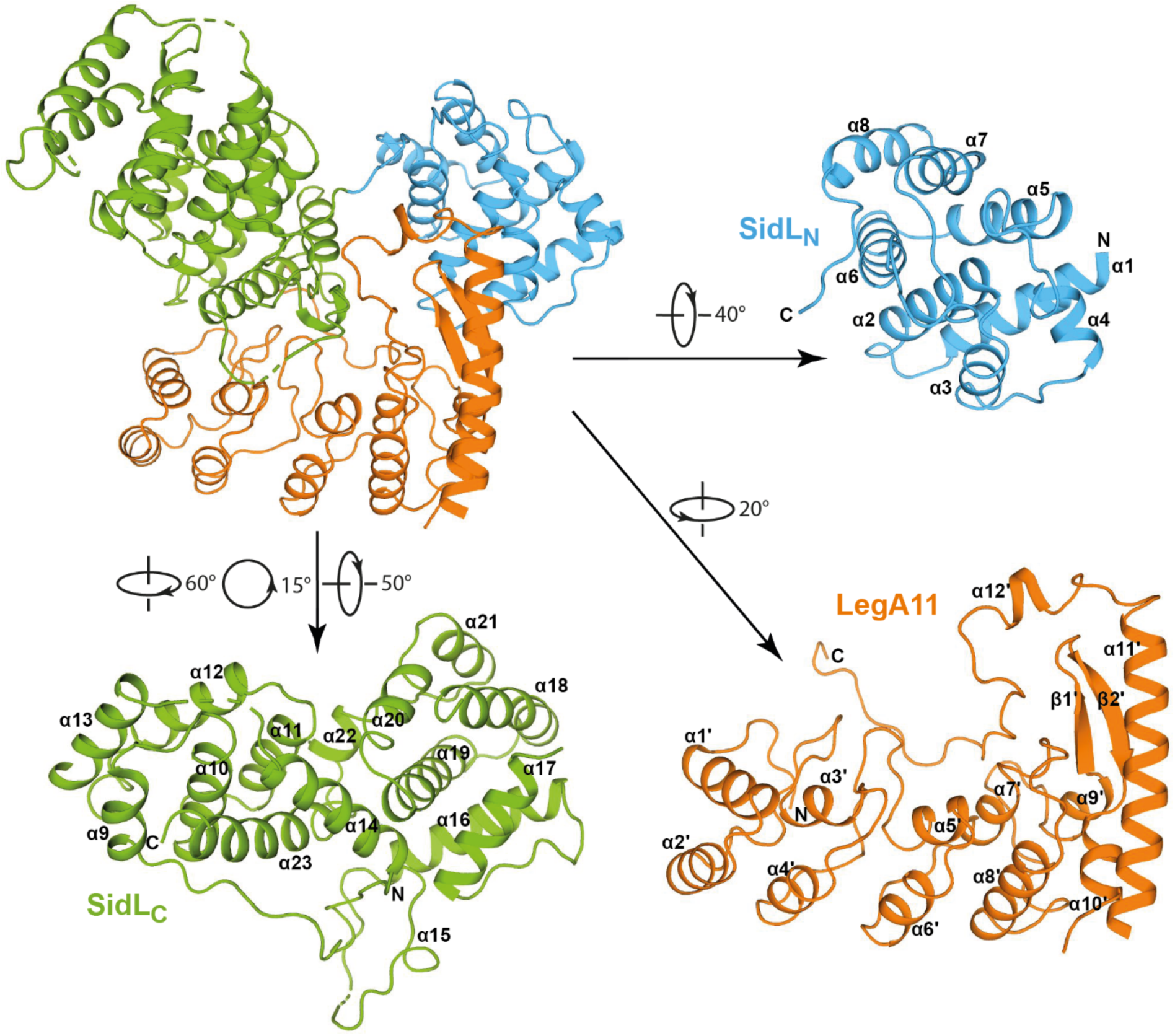
Overall structure of SidL/LegA11. Cartoon representation of the three-dimensional structure of the complex formed by SidL and LegA11. The model in the upper left corner represents the complex. The two subdomains of SidL are shown in blue and green, LegA11 is shown in orange. For better visibility, the complex components have been rotated individually, the respective degrees of rotation relative to the orientation of the complex are shown next to the arrows. Secondary structure elements as well as the N- and C-termini of the individual components are labeled.

SidLN consists of eight rather short helices (α1-α8) that form pairs, which are aligned in a nearly parallel fashion except for the second pair, where α4 is tilted by about 50° (Fig. 2). The order of the helix pairs is α3/α4, α1/α2, α5/α6, and α7/α8. Helices within pairs are connected via short loops, whereas the pairs are interconnected by longer loops. SidLC displays an elongated shape that is roughly divided into two parts. The first part contains helices α9-α13 that are arranged in parallel layers of three (α9-α11) and two (α12 and α13). The helix axes are oriented roughly parallel to each other. This part is completed by the C-terminal helix α23 that packs against the three-helix layer in an orthogonal fashion. The second part is composed of the helix pair α16 and α17 that packs orthogonally against a helix bundle formed by helices α18-α21. Both parts are connected via helix α14 that interacts with α23 and α19 and the short helix α22 preceding the terminal helix α23. α14 and α16 are connected via a long loop of 20 residues that contains the single-turn helix α15. Between α17 and α18, about 110 residues are missing that presumably have been removed by the protease used for crystallization.

The N-terminal part of LegA11 is formed by four ankyrin repeats, of which the first three display the typical arrangement of short, paired helices (α1’/α2’, α3’/α4’, α5’/α6’) with the connecting loops between pairs protruding at an angle of approximately 90° (Li et al., 2006). The fourth repeat (α7’/α8’) deviates from the other three through longer helices and a shortened loop between the third and fourth repeat. The first three repeats correspond to a previously published crystal structure (PDB ID 4ZHB) comprising residues 11-114. Both structures can be superimposed with a root mean square deviation of 0.52 Å. The fourth repeat is immediately followed by a pair of very short helices, α9’ and α10’. These helices are oriented roughly perpendicular to each other and are connected via a twisted two-stranded anti-parallel β-sheet formed by strands β1’ and β2’. The β-sheet is “wrapped” by the long helix α11’ following helix α10’ on one side, the single-turn helix α12’ at the top, and an extended loop on the other side. The overall shape resembles the letter L with the ankyrin repeats forming the stalk and the C-terminal extension forming the base.

A search for structural homologs using the DALI server with SidLN as search model did not yield significantly similar structures. An equivalent search using SidLC as search model identified the recently published cryo-EM structure of the *Legionella* effector LnaB (PDB ID 8JO4) as the closest structural homolog. Superposition of SidLC with the N-terminal domain of LnaB yielded an r.m.s.d. value of 2.7 Å for 201 equivalent Cα atoms (Suppl. Fig. 1). The identity of the aligned sequence stretches is 21.9 %.

### Characterization of the SidL/LegA11 binding interface

SidL interacts with LegA11 via an interface of about 2300 Å^2^ as determined by PISA and SPPIDER [30,31]. SidLC contacts LegA11 via the inside of both the stalk and the base of the L-shape with an interaction surface of about 1500 Å^2^, whereas SidLN is placed on the opposite side of the base of LegA11 (Fig. 2). The interaction of SidLN and LegA11 is mediated via several polar side chain-main chain interactions, for instance between the side chain of K180 of LegA11 with the main chain oxygen of R121 of SidL or the side chain of E118 of SidL with the main chain nitrogen of I178 of LegA11, respectively (Fig. 3). The only charged side chain-side chain interaction is between K224 of LegA11 and E118 of SidLN. Hydrophobic interactions involve residues L99 and L103 in SidLN and L232, I236, and F240 in LegA11. The interaction between SidLC and LegA11 is mainly mediated by polar and charged interactions, whereas distinct hydrophobic interactions are absent. In the following description, the respective residue of SidLC will be mentioned first. Salt bridges are formed between the side chains of D393 and of K86 and the side chains of R396 and of E254, respectively. Side chain-main chain interactions are visible between the side chain of D393 and the main chain nitrogen of E258, as well as of R401 and the carbonyl oxygens of F239 and T242. Main chain-main chain interactions exist between the carbonyl oxygen of K357 and the nitrogen of E245, and the carbonyl oxygen of F389 and the nitrogens of G261 and I262.

**Figure 3.**
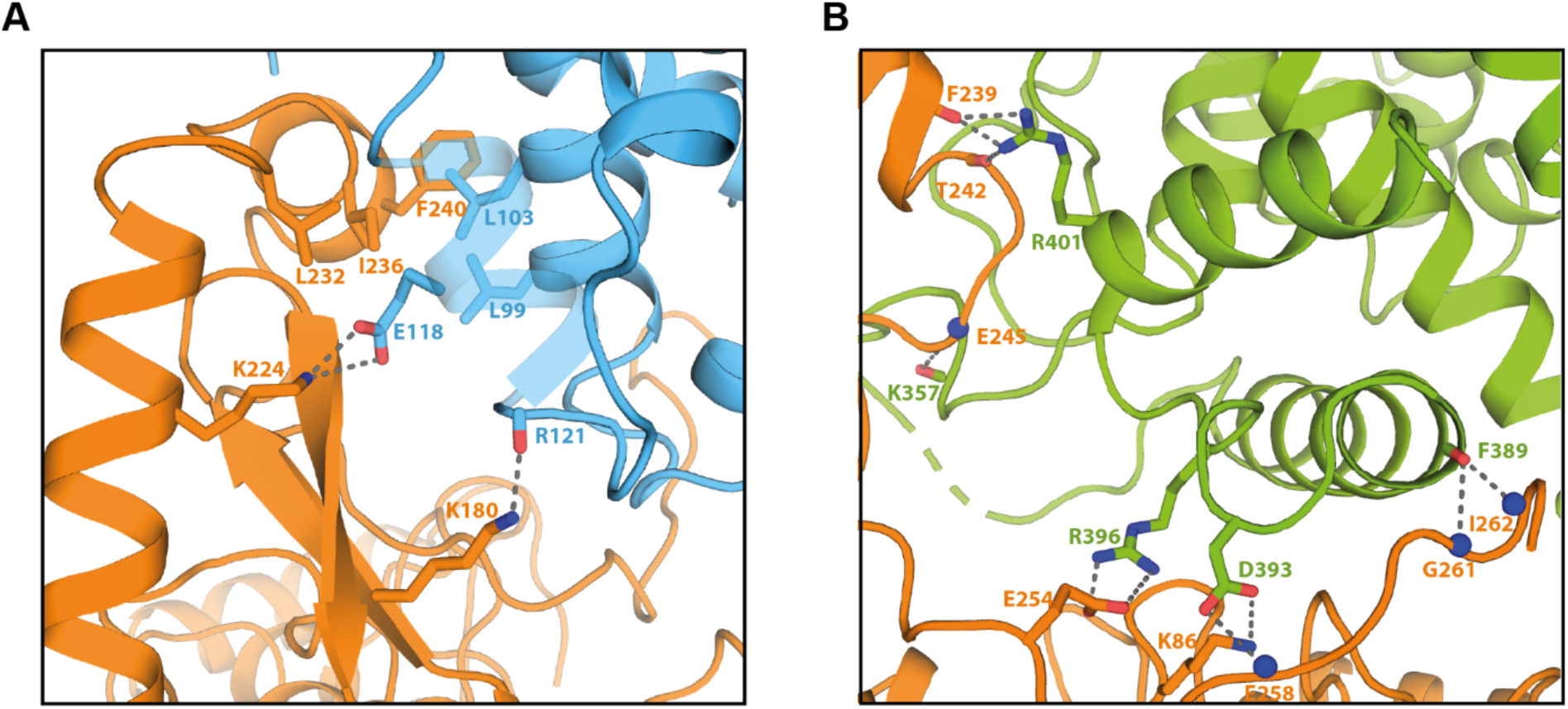
Binding interfaces of SidL and LegA11. Cartoon representation of the binding interfaces between SidL and LegA11. The color code corresponds to that in Figure 2. Side chains of amino acid residues responsible for mediating the interactions are depicted as stick models and are labeled accordingly. Hydrogen bonds and polar interaction are shown as grey dotted lines. **A.** Interface between SidLN and LegA11. **B.** Interface between SidLC and LegA11.

To determine the binding affinity of SidL and LegA11, we employed isothermal titration calorimetry (ITC) using purified proteins. Based on the domain boundaries of SidL seen in the crystal structure, we recombinantly produced the variant SidL76-628, but found the protein to be contaminated with a degradation product. Extending the C-terminus to residue K645 yielded a stable protein that could be purified with high purity and yield, which was used in subsequent ITC experiments. We performed ITC measurements with SidL in the sample cell at concentrations between 1.45 and 0.61 µM and a 20-fold molar excess of LegA11 in the syringe. Averaging the individual measurements, we obtained a dissociation constant of 1.8 nM, which is in good agreement with the stability of the SidL/LegA11 complex on the SEC column (Fig. 4).

**Figure 4.**
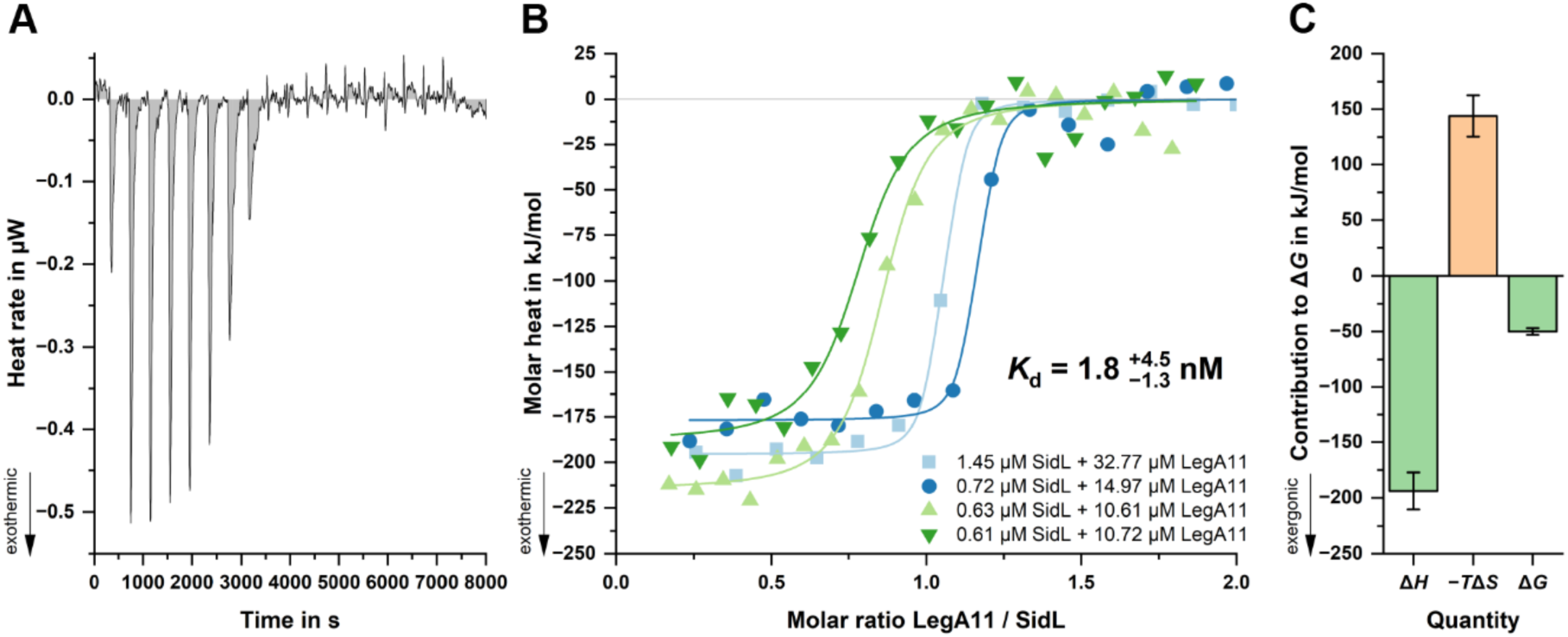
Affinity of SidL and LegA11 determined by ITC. **A.** Baseline-corrected heat rate (µW) versus time (s) for titration of LegA11 (ligand) into SidL (receptor) at 25 °C, showing exothermic binding events (negative peaks) that diminish after saturation. **B.** Integrated binding isotherms from four independent experiments plotted as molar heat (kJ/mol) versus molar ratio of LegA11 to SidL. The best-fit one-site model yielded a dissociation constant *K*d of 1.8 ^+4.5^ −1.3 nM, with a stoichiometry of 0.92 ± 0.16. Binding was strongly exothermic (average Δ*H* ≈ −200 kJ/mol). Receptor and ligand concentrations were: 1.45/32.77 µM, 0.72/14.97 µM, 0.63/10.61 µM, and 0.61/10.72 µM (SidL/LegA11). **C.** Thermodynamic signature of binding showing large enthalpic contribution (Δ*H* = −194 ± 17 kJ/mol) opposed by an unfavorable entropic term (−*T*Δ*S* = +144 ± 16 kJ/mol), resulting in a free energy change of Δ*G* = −50 ± 3 kJ/mol.

To verify the binding interface between SidL and LegA11 seen in the crystal structure, we introduced mutations into LegA11 to disturb complex formation. Given the large size of the interface, we decided to pursue a rather drastic approach by generating the quadruple mutant K95A/F239R/F240R/G243R and purified the resulting protein (LegA11-4M) to homogeneity. We subjected SidL76-645 alone, LegA11 in wild-type or mutant form alone, or in equimolar ratio with SidL76-645 to analytical size exclusion chromatography (SEC). As expected, wild-type LegA11 formed a stable complex with the truncated SidL variant (Fig. 5). The chromatogram corresponding to SidL76-645 and LegA11-4M shows two distinct peaks very close to the positions of the single proteins. The slight shift to lower elution volumes most likely reflects a very weak transient interaction between SidL76-645 and LegA11-4M. We conclude that the interface observed in the crystal structure is indeed responsible for binding of LegA11 to SidL and that the interaction is largely abrogated by introduction of the mutations.

**Figure 5.**
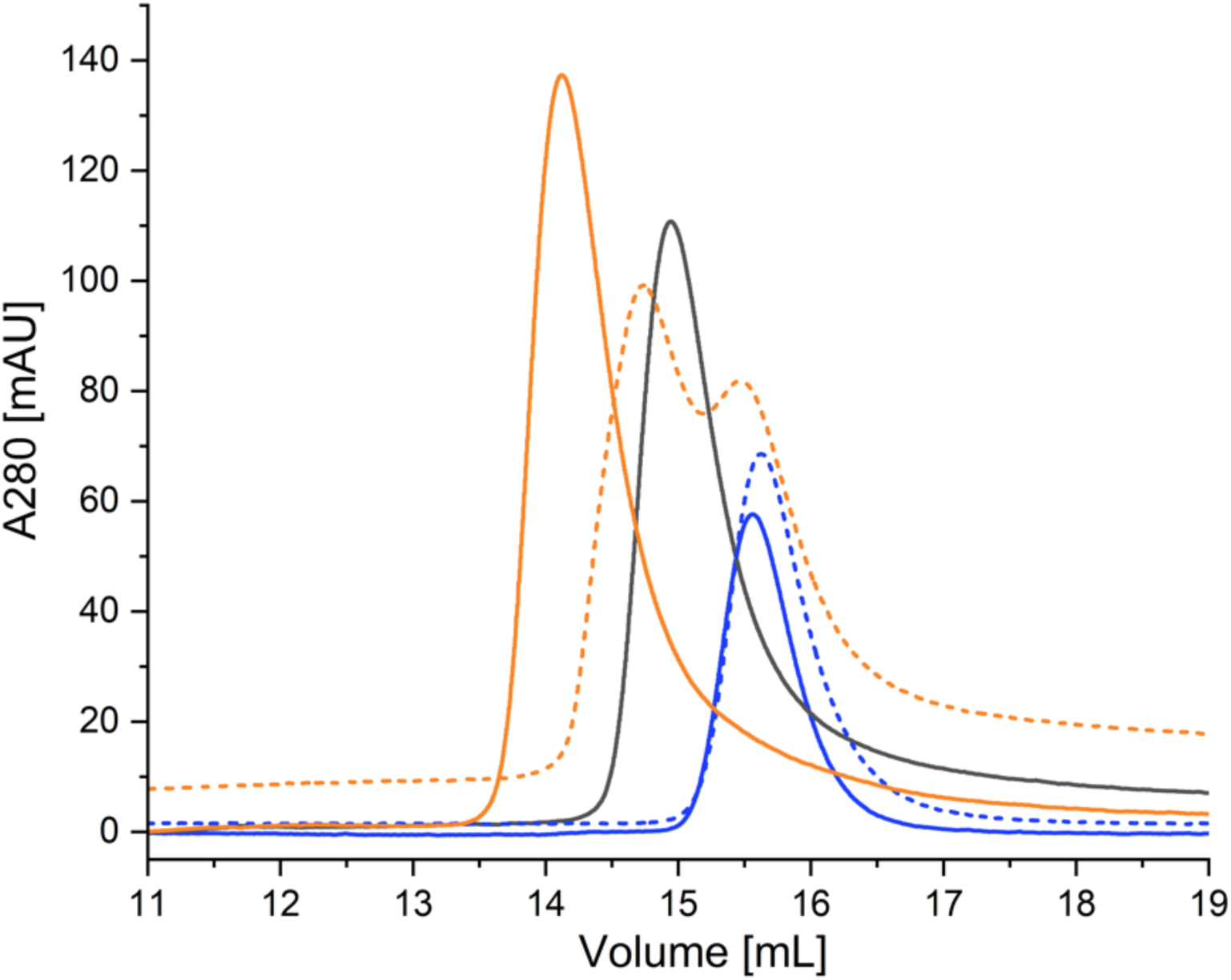
Influence of mutations in LegA11 on binding to SidL76-645. 1 mg SidL76-645 and/or 400 µg LegA11 in 100 µL buffer were applied to an S200 10/300 increase size exclusion column. The colors of the respective chromatograms are as follows: SidL76-645 – black; LegA11-WT – blue; LegA11-4M – dotted blue; SidL76-645/LegA11-WT – orange; SidL76-645/LegA11-4M – dotted orange.

### LegA11 restores SidL-inhibited mRNA translation via a direct protein-protein interaction

Of the eight *L. pneumophila* effectors that block eukaryotic mRNA translation, two are associated with metaeffectors. The translation inhibiting effects of SidI are completely abolished in the presence of its metaeffector, MesI (Joseph et al., 2020). Therefore, we evaluated the effects of LegA11 on SidL-mediated translation inhibition. Recombinant His6-Myc-SidL and -LegA11 were purified from *E. coli* (Suppl. Fig 2) and added to cell-free rabbit reticulocyte lysates to quantify luciferase (Luc) mRNA translation. SidL inhibited Luc mRNA translation in a dose-dependent manner but with less potency than the effector SidI (Fig 6A), validating results from a prior study [9]. LegA11 binds and abrogates SidL toxicity in a heterologous yeast expression system [14]; thus, we tested the hypothesis that LegA11 suppresses SidL-mediated translation inhibition. Translation increased significantly when equimolar amounts of LegA11 and SidL were added to reactions and did not differ from reactions containing LegA11 alone (Fig 6B). Thus, LegA11 binds and suppresses SidL activity, further validating its role as a metaeffector of SidL.

**Figure 6.**
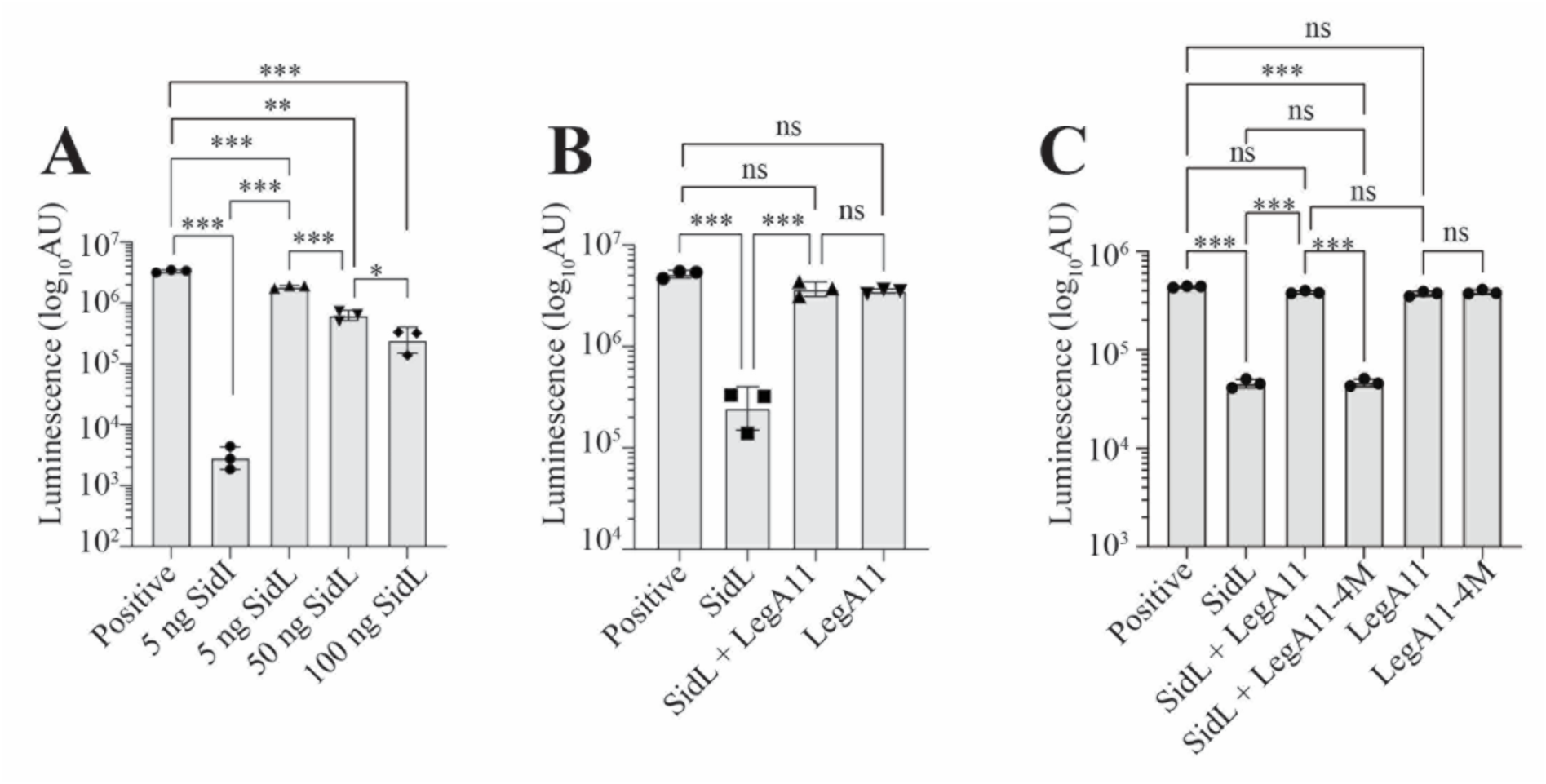
LegA11 restores SidL-mediated translation inhibition via a direct protein-protein interaction. Rabbit reticulocyte lysates were incubated with Firefly luciferase (Luc) mRNA alone (Positive) or with (A) 5 ng SidI, 5ng SidL, 50 ng SidL, or 100 ng SidL; (B) 65 nM of SidL, SidL and LegA11, or LegA11 alone; or (C) 65 nM of SidL alone, SidL and LegA11 variants, or LegA11 variants alone. Data shown are mean ± standard deviation (s.d.) of samples in triplicates (N=3) and are representative of three independent experiments. AU, arbitrary units. Asterisks (*) denote statistical significance (\*\*\**P*<0.001, \**P*<0.05, ns, not significant) by one-way ANOVA with Tukey’s post-hoc test.

We hypothesized that SidL-LegA11 complex formation is required for functional regulation of SidL. To test this, we quantified Luc mRNA translation in cell-free lysates containing equimolar amounts of purified recombinant SidL with either wild-type LegA11 or LegA11-4M (Suppl. Fig. 2), the latter of which has lost the ability to bind SidL (Fig 5). SidL-mediated translation inhibition was suppressed by wild-type LegA11 but unaffected by LegA11-4M since the extent of translation inhibition did not differ from reactions containing SidL alone (Fig. 6C). Luc mRNA translation was not affected by either wild-type LegA11 or LegA11-4M alone (Fig. 6C). Together, these data suggest that LegA11 regulates SidL via a direct protein-protein interaction and that SidL is unable to inhibit translation when it is bound to LegA11.

Neither SidL nor LegA11 impact L. pneumophila replication.

He *et al.* found that neither SidL nor LegA11 individually influence *L. pneumophila* replication within primary bone marrow-derived macrophages (BMDMs) from susceptible A/J mice and the social amoebae, *Dictyostelium discoideum* [15]. *D. discoideum* is routinely used as a model to study *L. pneumophila* in the laboratory; however, it is not naturally found in freshwater environments colonized by *L. pneumophila* [32,33]. To determine whether SidL or LegA11 influence *L. pneumophila* replication within a natural protozoan host, *L. pneumophila* strains with clean unmarked chromosomal deletions of *legA11* or *sidL* (Δ*legA11* or Δ*sidL*, respectively) were generated to assess replication within *Acanthamoeba castellanii.* We found no differences in *L. pneumophila* Δ*legA11* or Δ*sidL* growth relative to either wild-type *L. pneumophila* or genetically complemented strains in *A. castellanii* (Suppl. Fig 3A). We also found no differences in growth in AYE broth *in vitro* (Suppl. Fig. 3B). These data support prior data suggesting that SidL and LegA11, like many *L. pneumophila* effectors, function redundantly with other effectors in *A. castellanii* [20,34].

## Discussion

Over the past decade, a growing number of metaeffectors have been identified and functionally characterized [10,12–14,20,35]. Metaeffectors are defined as effectors targeting another effector by direct physical interactions [11], as opposed to a biochemical activity that antagonizes the activity of a different effector [36–38]. LegA11 had been previously characterized as a putative metaeffector of SidL [14,15], but the role of the SidL-LegA11 interaction in LegA11’s metaeffector activity was unknown. Using purified proteins, we found that SidL and LegA11 bind with high affinity to form a stable binary complex. This finding prompted us to attempt a crystallographic characterization of the complex. Despite extensive crystallization trials using different N- and C-terminal truncation variants of SidL, we only obtained crystals after adding trace amounts of protease. This suggests that SidL contains one or more flexible regions that counteract crystal formation and is in line with the observation that SidL in the crystal structure is further truncated both N- and C-terminally than our shortest construct used for crystallization. Furthermore, a portion of 110 amino acid residues (aa 414-524) had been removed from SidLC opposite the LegA11 binding site, which may indicate that this portion is flexible in the absence of a putative second binding partner.

SidL and LegA11 interact via a rather large interface of about 2300 Å^2^ involving mostly charged or polar side chains (Fig. 2 + 3). Although such an interface size suggests biological relevance [39], we wanted to verify that this interface is not a crystallization artifact. Indeed, mutagenesis of four interface residues in LegA11 (LegA11-4M) disrupted the interaction between the two proteins) We found that LegA11-4M was unable to form a complex with SidL (Fig 5), confirming the validity of the interface seen in the crystal structure (Fig. 3). The inability of LegA11-4M to suppress SidL’s translation inhibition (Fig 6C) indicates that LegA11’s metaeffector activity is mediated by a direct protein-protein interaction with SidL.

A very recent study describes SidL as an ATPase that hydrolyzes ATP to AMP and pyrophosphate (He et al., 2025). SidL binds ATP via an S-HxxxE motif [15], which is also present in toxins from phylogenetically diverse bacterial species [40]. He et al. (2025) suggested that the inhibitory effect of LegA11 is exerted via binding to the C-terminal domain of SidL, where the active center is located [15]. To examine the spatial relationship between LegA11 and the ATP-binding site, we generated an AlphaFold3 (AF3) model of SidL complexed with ATP and superposed it with the SidL from our crystal structure. From the superposition it is apparent that LegA11 does not bind in the vicinity of the active site (Suppl. Fig. 4). Thus, it is unlikely that LegA11 abolishes ATP hydrolysis by directly occluding the nucleotide binding site of SidL. He et al. suggest that SidL functions to deplete cytosolic ATP pools in the host cell cytosol, which could plausibly limit mRNA translation; however, it is also tempting to speculate a mechanistic connection between impaired actin polymerization and translation since F-actin serves as a scaffold for polyribosome assembly [9,41–45]. Sustained translation on monoribosomes may explain why SidL blocks translation with less potency than the effector SidI, which directly inactivates translating ribosomes.

SidL is one of at least eight *Legionella* effectors able to inhibit host mRNA translation *in vitro* [46–48]. The translation-inhibiting effector SidI (Lpg2504) is also regulated by a metaeffector, MesI (<u>Me</u>taeffector of <u>Si</u>dI; Lpg2505). Like SidL and LegA11, SidI and MesI also form a high-affinity 1:1 complex that renders SidI non-toxic and unable to suppress mRNA translation [12]. Despite functional similarities, SidL and SidI are mechanistically distinct since SidI is a mannosyltransferase that directly inactivates the host mRNA translation machinery [9,12]. Moreover, unlike for SidL, the enzymatic activity of SidI is independent of host co-factors and requires only its activated sugar donor, GDP-mannose, for catalysis [12]. While SidI can function within *L. pneumophila* by itself [49], the requirement for host actin as a co-factor provides a plausible explanation for lack of intrabacterial SidL activity (Suppl. Fig 3B).

SidL’s requirement for globular actin as an activating co-factor is similar to actin-mediated activation of the *Legionella* effector LnaB [15,40,50]. He *et al.* found that SidL binds LegA11 and actin simultaneously [15], which we validated using AF3 to generate a hypothetical ternary complex by overlaying SidL from our crystal structure with the SidL from the actin complex (Suppl. Fig. 5). Indeed, the binding sites of LegA11 and actin do not overlap, confirming the experimental findings of He et al. Interestingly, in all 15 AF3 predictions, SidLC is sufficient for binding actin, whereas SidLN is able to assume a wide variety of conformations (Suppl. Fig. 6). This is in line with the binding mode of actin in several crystal structures with LnaB, which does not possess a subdomain corresponding to SidLN. It is evident from our crystal structure that upon binding, LegA11 abolishes the conformational flexibility of SidLN and locks it in a fixed orientation relative to SidLC. This conformational restriction is also likely reflected in the unusually high loss of entropy (−*T*Δ*S* = 144 kJ/mol) seen in our ITC measurements (Fig. 4).

A recent structural study showed that actin enhances the affinity of LnaB for its target phosphoribosyl-ubiquitin (PR-Ub) [51]. According to these authors’ model, the hydrolysis incompetent binary complex formed by LnaB and actin is activated by binding of PR-Ub. However, for SidL no third protein is required, since purified SidL and actin are sufficient for efficient ATP hydrolysis [15]. Therefore, actin must play a direct role in inducing hydrolytic competence. The most likely scenario is that binding of actin orders the large flexible loop (residues 414-524) of SidL, which at its C-terminal end borders on the nucleotide binding site. This reorientation may position loop residues required for nucleotide hydrolysis in a catalytically competent fashion. We hypothesize that the fixed N-terminal domain seen in our crystal structure prevents the correct positioning of the loop, effectively rendering SidL catalytically inactive despite concomitant binding of actin. However, elucidation of the precise mechanisms of both activation by actin and inhibition by LegA11 will require structural characterization of the respective complexes in the future.

## Acknowledgments

We thank Petra Hinse for excellent technical assistance, Dr. Craig Roy for providing pSN85 and pSR47S plasmids, and Dr. Brian Geisbrecht (Kansas State University) for helpful discussions and providing the pT7HMT plasmid.

## Disclosure Statement

No potential conflict of interest was declared by the authors.

## Funding

Work in the Shames lab was supported by the National Institutes of Health (P20GM113117) and start-up funds from Kansas State University and Michigan State University.

## Author contributions

Conceptualization, S.R.S., T.F.R.; methodology, investigation, validation and data curation, D.A.M., C.A.H., A.M.S., J.E., J.M.W., S.E., S.R.S., and T.F.R.; writing – original draft preparation, S.R.S. and T.F.R.; writing – review and editing, D.A.M., C.A.H., A.M.S., J.E., J.M.W., S.E., S.R.S., and T.F.R.; supervision, S.E., S.R.S., and T.F.R.; All authors have read and agreed to the published version of the manuscript.

## Supplemental Information

**Suppl. Table 1.**
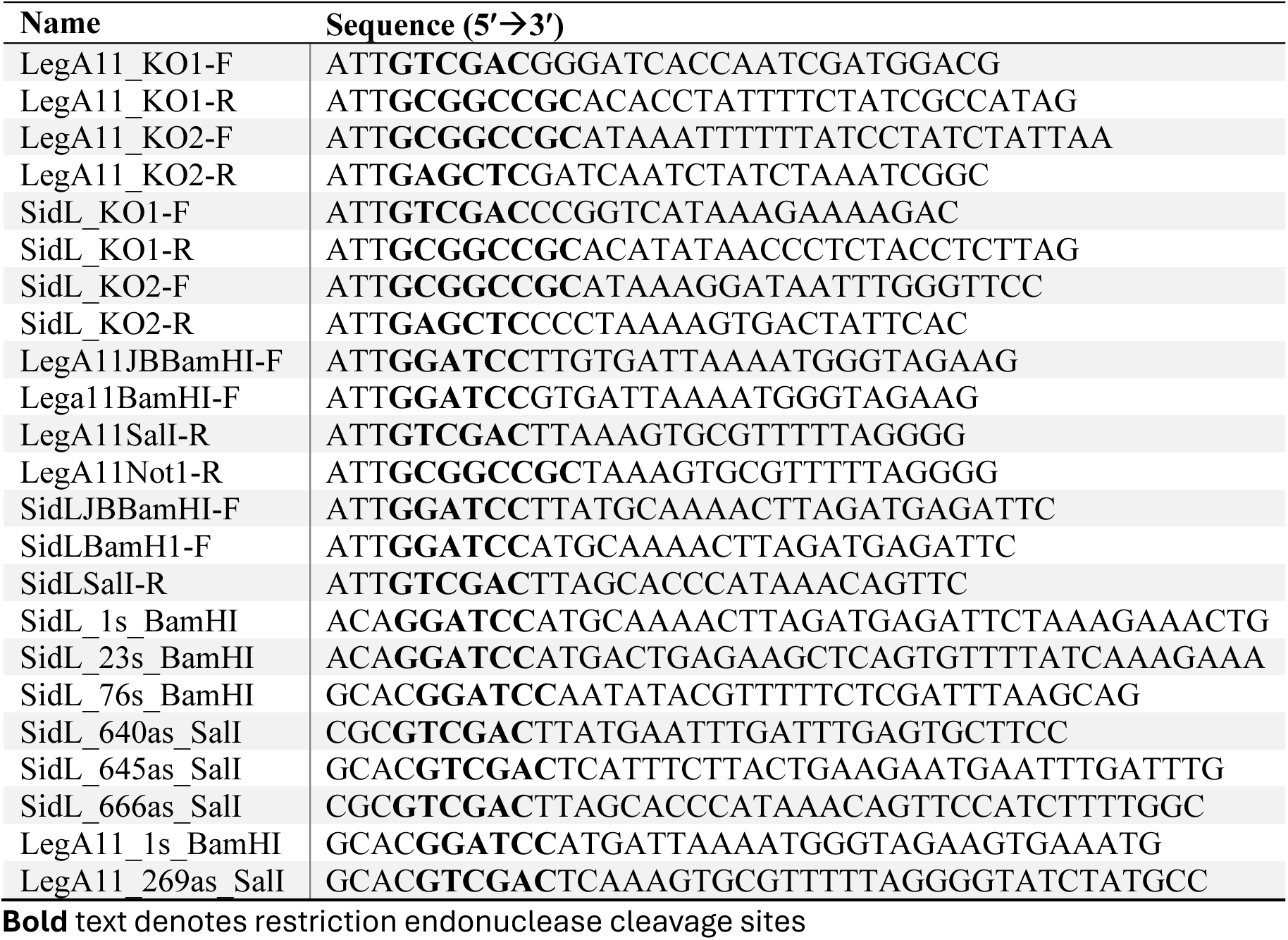
Oligonucleotide primers used in this study.

**Suppl. Fig. 1.**
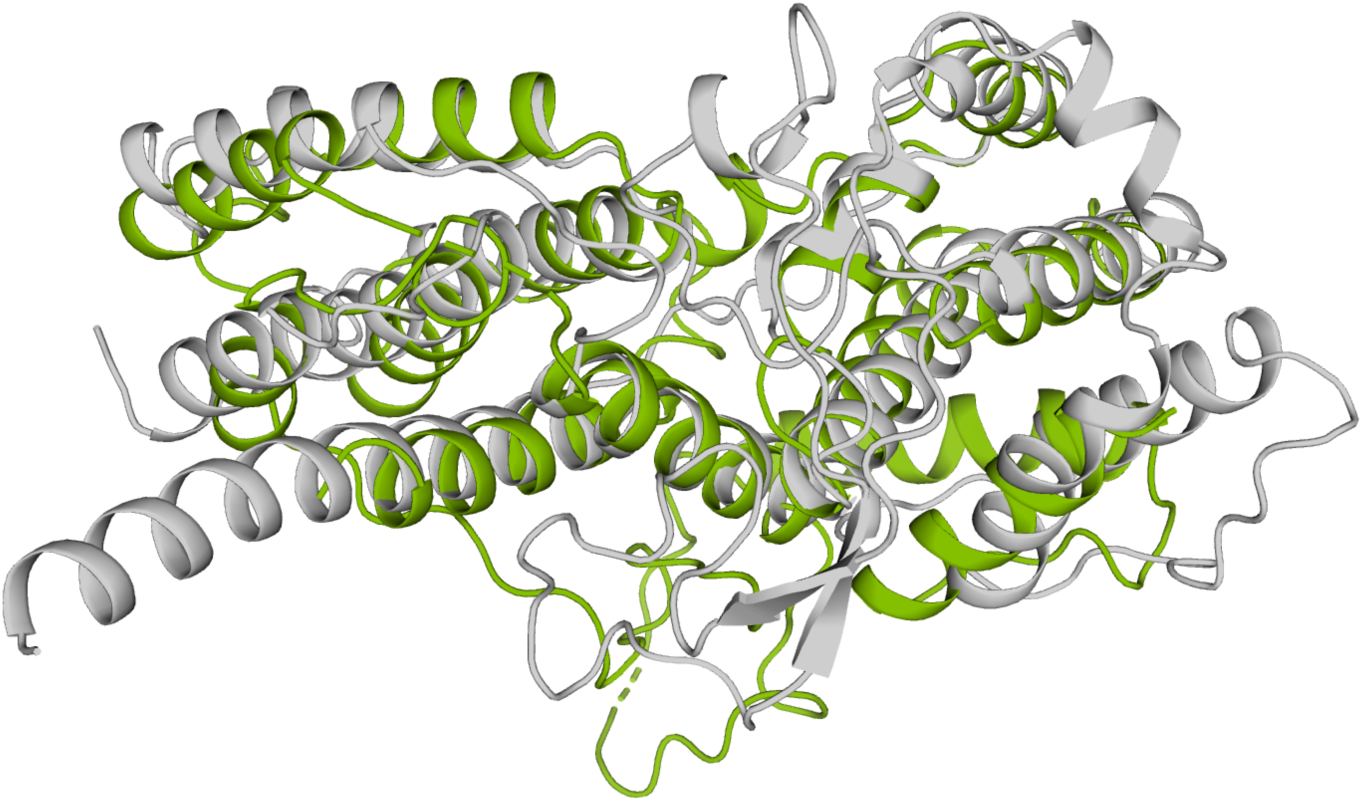
Superposition of SidL and LnaB. Cartoon representation of the three-dimensional structure of SidLC superposed to a crystal structure of LnaB of *L. pneumophila* (PDB ID 8JO3). SidL is shown in green, LnaB is shown in gray.

**Suppl. Fig 2.**
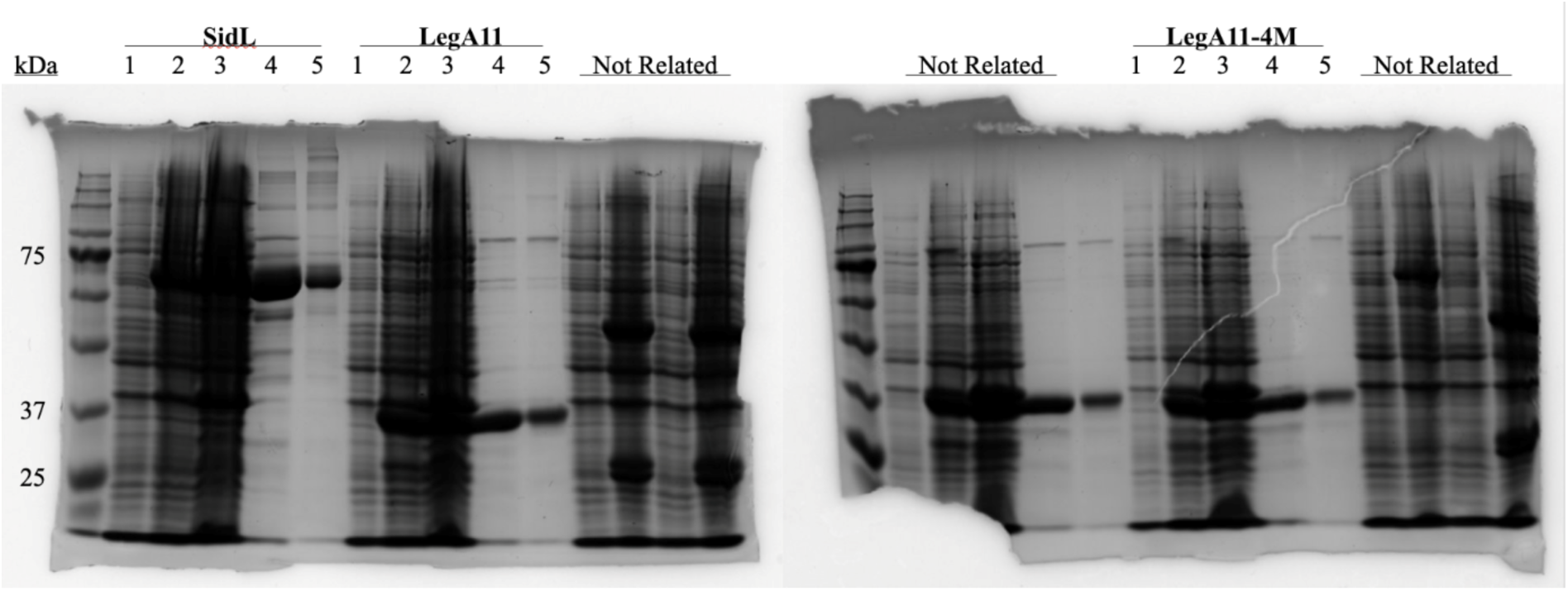
Purification of recombinant SidL and LegA11 for in vitro translation assays. Purification of recombinant SidL and LegAll variants was visualized via SDS-Page and Coomassie Brilliant Blue staining. Proteins for each set of lanes denoted above. Lanes numbered as follows: 1) Uninduced 2) Induced 3) Sonicated 4) Purified protein 5) Dialysed purified protein.

**Suppl. Fig. 3.**
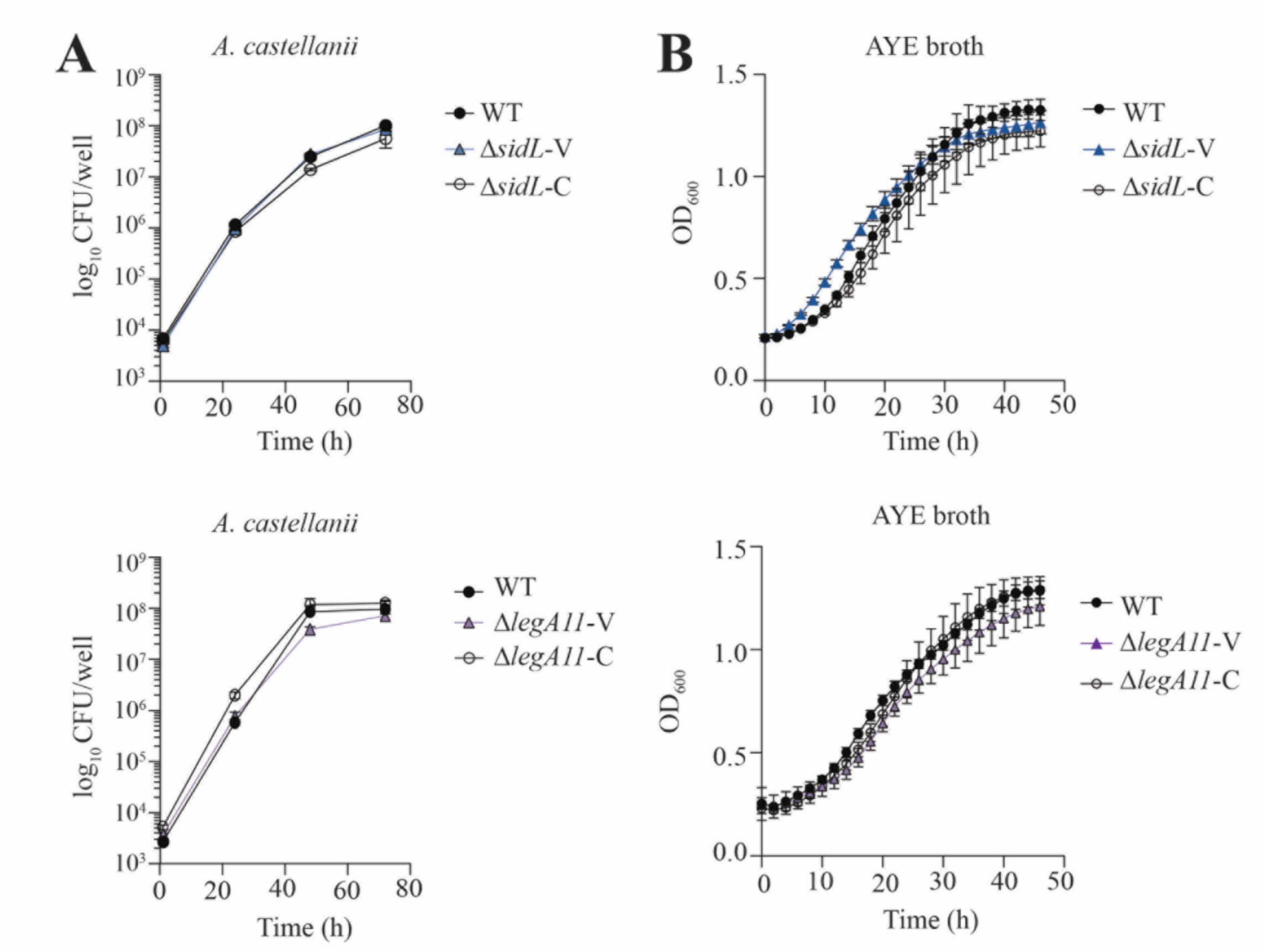
Neither SidL nor LegA11 impact *L. pneumophila* intracellular replication or growth *in vitro*. **(A)** Colony forming units (CFU) recovered from *Acanthamoeba castellanii* infected with the indicated *L. pneumophila* strains (MOI of 0.1) for the indicated times. **(B)** Optical density (OD) 600 nm values of *L. pneumophila* grown in AYE broth. Data were collected every 2h for up to 46h and shown as mean ± standard deviation (s.d.) of six individual wells (N=6). Representative of two independent experiments. Data shown are mean ± s.d. of three independent wells (N=3) and representative of three independent experiments. V denotes empty plasmid vector, and C denotes plasmid-based genetic complementation construct.

**Suppl. Fig. 4.**
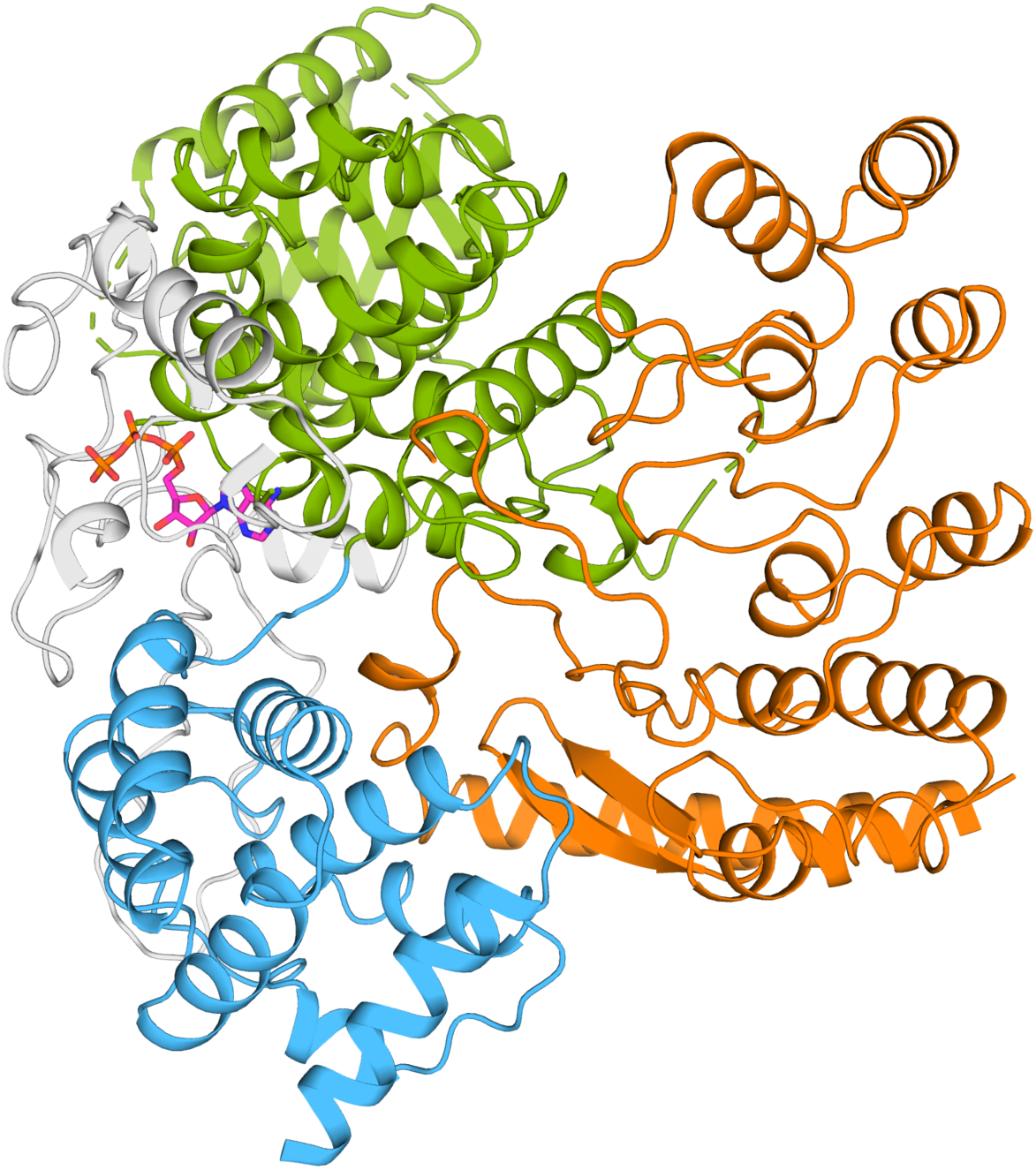
Spatial relationship between the binding sites of LegA11 and ATP in SidL. Cartoon representation of the crystal structure of the SidL/LegA11 complex. The color code corresponds to that in Fig. 2. The ATP molecule from an AF3 prediction of SidL complexed with ATP is shown as stick model. The coloring of the ATP is as follows: carbon – magenta; nitrogen – blue; oxygen – red; phosphorus – orange.

**Suppl. Fig. 5.**
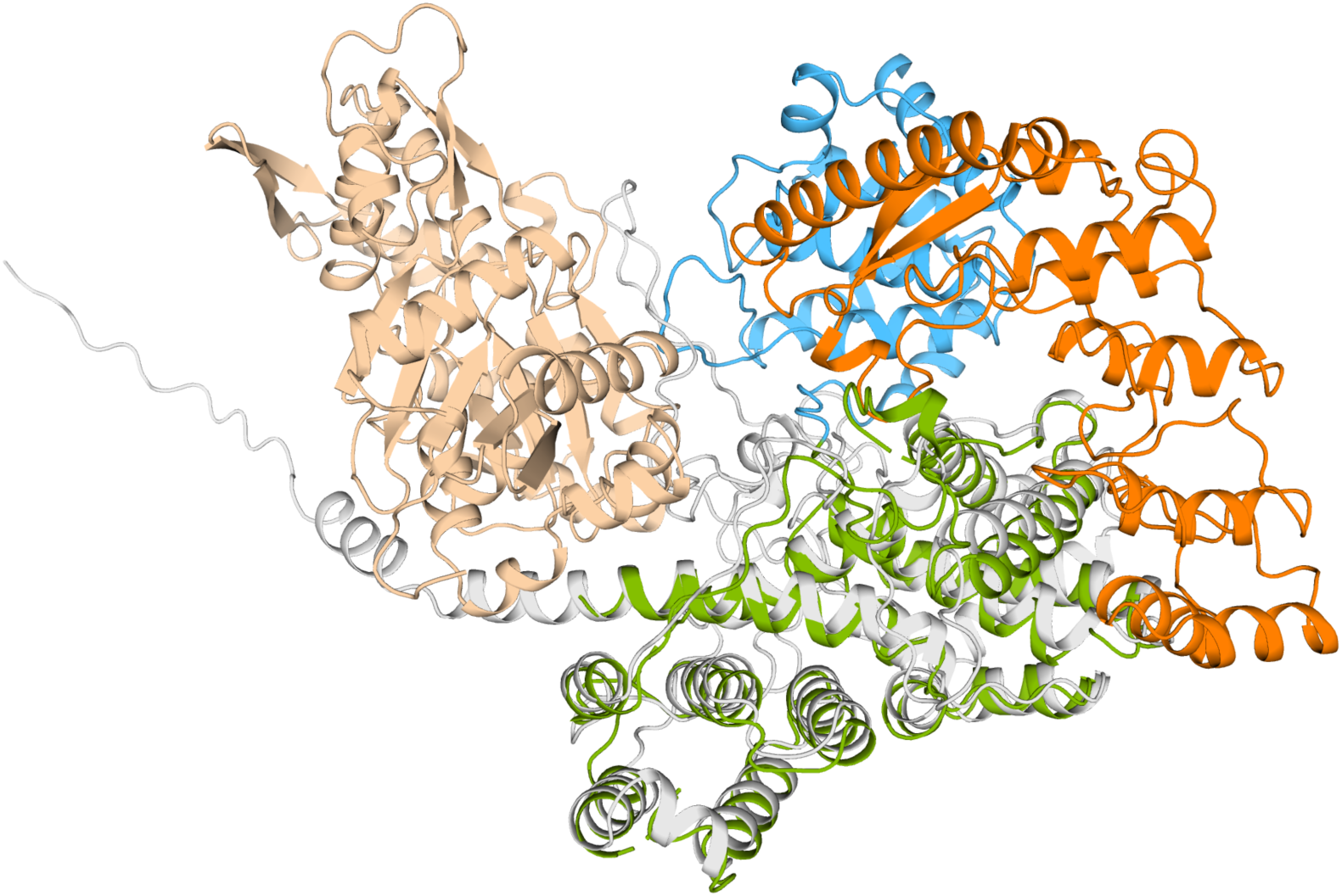
Hypothetical ternary complex of SidL, LegA11, and actin. SidL from the crystal structure was superposed with SidLC from an AF3 prediction of SidL and actin. Components of the crystal structure are colored as in Fig. 2, SidLC from the AF3 prediction is shown in gray, actin is shown in beige.

**Suppl. Fig. 6.**
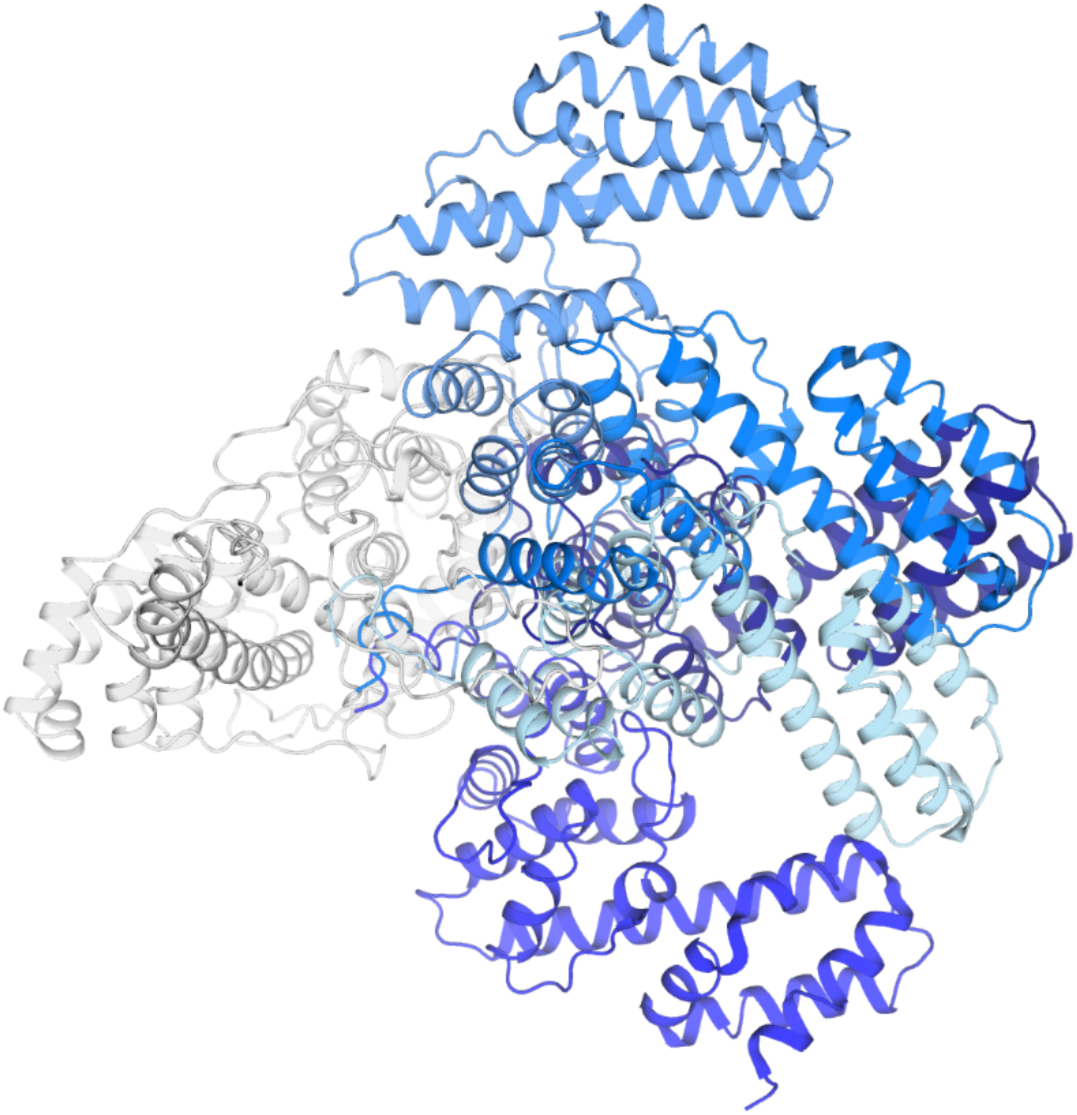
Conformational flexibility of SidLN in AF3 predictions of complexes between SidL and actin. Cartoon representation of a superposition of the SidL molecules of five AF3 predictions of a SidL/actin complex. The models are superposed via SidLC. SidLN of the different models are shown in different shades of blue.

## Notes

### Competing Interest Statement

The authors have declared no competing interest.

## References

1. Gomez-Valero L, Buchrieser C. Intracellular parasitism, the driving force of evolution of Legionella pneumophila and the genus Legionella. Microbes Infect. 2019;21: 230–236. doi:10.1016/j.micinf.2019.06.012

2. Lockwood DC, Amin H, Costa TRD, Schroeder GN. The Legionella pneumophila Dot/Icm type IV secretion system and its effectors. Microbiology (Reading). 2022;168. doi:10.1099/mic.0.001187

3. Mondino S, Schmidt S, Rolando M, Escoll P, Gomez-Valero L, Buchrieser C. Legionnaires’ Disease: State of the Art Knowledge of Pathogenesis Mechanisms of Legionella. Annu Rev Pathol. 2020;15: 439–466. doi:10.1146/annurev-pathmechdis-012419-032742

4. Romanov KA, O’Connor TJ. Legionella pneumophila, a Rosetta stone to understanding bacterial pathogenesis. J Bacteriol. 2024;206: e0032424. doi:10.1128/jb.00324-24

5. Belyi Y, Jank T, Aktories K. Effector glycosyltransferases in legionella. Front Microbiol. 2011;2: 76. doi:10.3389/fmicb.2011.00076 [doi]

6. Belyi Y, Tabakova I, Stahl M, Aktories K. Lgt: a family of cytotoxic glucosyltransferases produced by Legionella pneumophila. J Bacteriol. 2008;190: 3026–35. doi:10.1128/JB.01798-07

7. Moss SM, Taylor IR, Ruggero D, Gestwicki JE, Shokat KM, Mukherjee S. A Legionella pneumophila Kinase Phosphorylates the Hsp70 Chaperone Family to Inhibit Eukaryotic Protein Synthesis. Cell Host Microbe. 2019;25: 454–462.e6. doi:S1931-3128(19)30043-5 [pii]

8. Syriste L, Patel DT, Stogios PJ, Skarina T, Patel D, Savchenko A. An acetyltransferase effector conserved across Legionella species targets the eukaryotic eIF3 complex to modulate protein translation. mBio. 2024;15: e0322123. doi:10.1128/mbio.03221-23

9. Subramanian A, Wang L, Moss T, Voorhies M, Sangwan S, Stevenson E, et al. A Legionella toxin exhibits tRNA mimicry and glycosyl transferase activity to target the translation machinery and trigger a ribotoxic stress response. Nat Cell Biol. 2023;25: 1600–1615. doi:10.1038/s41556-023-01248-z

10. Kubori T, Shinzawa N, Kanuka H, Nagai H. Legionella metaeffector exploits host proteasome to temporally regulate cognate effector. PLoS Pathog. 2010;6: e1001216. doi:10.1371/journal.ppat.1001216 [doi]

11. Joseph AM, Shames SR. Affecting the Effectors: Regulation of Legionella pneumophila Effector Function by Metaeffectors. Pathogens. 2021;10. doi:10.3390/pathogens10020108

12. Joseph AM, Pohl AE, Ball TJ, Abram TG, Johnson DK, Geisbrecht B V, et al. The Legionella pneumophila metaeffector Lpg2505 (MesI) regulates SidI-mediated translation inhibition and novel glycosyl hydrolase activity. Infect Immun. 2020. doi:IAI.00853-19 [pii]

13. Machtens DA, Willerding JM, Eschenburg S, Reubold TF. Crystal structure of the metaeffector MesI (Lpg2505) from Legionella pneumophila. Biochem Biophys Res Commun. 2020;527: 696–701. doi:10.1016/j.bbrc.2020.05.027

14. Urbanus ML, Quaile AT, Stogios PJ, Morar M, Rao C, Leo R Di, et al. Diverse mechanisms of metaeffector activity in an intracellular bacterial pathogen, Legionella pneumophila. Mol Syst Biol. 2016;12: 893. doi:10.15252/msb.20167381 [doi]

15. He C, Li C, Liu Y, Chen T-T, Li C, Chu X, et al. Modulation of host ATP levels by secreted bacterial effectors. Nat Commun. 2025;16: 4675. doi:10.1038/s41467-025-60046-3

16. Feeley JC, Gibson RJ, Gorman GW, Langford NC, Rasheed JK, Mackel DC, et al. Charcoal-yeast extract agar: primary isolation medium for Legionella pneumophila. J Clin Microbiol. 1979;10: 437–41. doi:10.1128/jcm.10.4.437-441.1979

17. Moffat JF, Tompkins LS. A quantitative model of intracellular growth of Legionella pneumophila in Acanthamoeba castellanii. Infect Immun. 1992;60: 296–301. doi:10.1128/iai.60.1.296-301.1992

18. Geisbrecht B V, Bouyain S, Pop M. An optimized system for expression and purification of secreted bacterial proteins. Protein Expr Purif. 2006;46: 23–32. doi:10.1016/j.pep.2005.09.003

19. Folly-Klan M, Alix E, Stalder D, Ray P, Duarte L V, Delprato A, et al. A novel membrane sensor controls the localization and ArfGEF activity of bacterial RalF. PLoS Pathog. 2013;9: e1003747. doi:10.1371/journal.ppat.1003747

20. Shames SR, Liu L, Havey JC, Schofield WB, Goodman AL, Roy CR. Multiple Legionella pneumophila effector virulence phenotypes revealed through high-throughput analysis of targeted mutant libraries. Proc Natl Acad Sci U S A. 2017;114: E10446–E10454. doi:10.1073/pnas.1708553114 [doi]

21. Nagai H, Roy CR. The DotA protein from Legionella pneumophila is secreted by a novel process that requires the Dot/Icm transporter. EMBO J. 2001;20: 5962–70. doi:10.1093/emboj/20.21.5962

22. Machtens DA, Willerding JM, Eschenburg S, Reubold TF. Crystal structure of the N-terminal domain of the effector protein SidI of Legionella pneumophila reveals a glucosyl transferase domain. Biochem Biophys Res Commun. 2023;661: 50–55. doi:10.1016/j.bbrc.2023.04.029

23. Studier FW. Protein production by auto-induction in high density shaking cultures. Protein Expr Purif. 2005;41: 207–34. doi:10.1016/j.pep.2005.01.016

24. Kabsch W. XDS. Acta Crystallogr D Biol Crystallogr. 2010;66: 125–132.

25. Kabsch W. Integration, scaling, space-group assignment and post-refinement. Acta Crystallogr D Biol Crystallogr. 2010;66: 133–144.

26. Skubak P, Pannu NS. Automatic protein structure solution from weak X-ray data. Nat Commun. 2013;4: 2777. doi:10.1038/ncomms3777 [doi]

27. Liebschner D, Afonine P V, Baker ML, Bunkoczi G, Chen VB, Croll TI, et al. Macromolecular structure determination using X-rays, neutrons and electrons: recent developments in Phenix. Acta crystallographicaSection D, Structural biology. 2019;75: 861–877. doi:10.1107/S2059798319011471 [doi]

28. Emsley P, Lohkamp B, Scott WG, Cowtan K. Features and development of Coot. Acta Crystallogr D Biol Crystallogr. 2010;66: 486–501.

29. Wernimont A, Edwards A. In situ proteolysis to generate crystals for structure determination: an update. PLoS One. 2009;4: e5094. doi:10.1371/journal.pone.0005094

30. Krissinel E, Henrick K. Inference of macromolecular assemblies from crystalline state. J Mol Biol. 2007;372: 774–97. doi:10.1016/j.jmb.2007.05.022

31. Langdon QK, Peris D, Kyle B, Hittinger CT. sppIDer: A Species Identification Tool to Investigate Hybrid Genomes with High-Throughput Sequencing. Mol Biol Evol. 2018;35: 2835–2849. doi:10.1093/molbev/msy166

32. Solomon JM, Isberg RR. Growth of Legionella pneumophila in Dictyostelium discoideum: a novel system for genetic analysis of host-pathogen interactions. Trends Microbiol. 2000;8: 478–80. doi:10.1016/s0966-842x(00)01852-7

33. Hägele S, Köhler R, Merkert H, Schleicher M, Hacker J, Steinert M. Dictyostelium discoideum: a new host model system for intracellular pathogens of the genus Legionella. Cell Microbiol. 2000;2: 165–71. doi:10.1046/j.1462-5822.2000.00044.x

34. Park JM, Ghosh S, O’Connor TJ. Combinatorial selection in amoebal hosts drives the evolution of the human pathogen Legionella pneumophila. Nat Microbiol. 2020;5: 599– 609. doi:10.1038/s41564-019-0663-7

35. Black MH, Osinski A, Gradowski M, Servage KA, Pawłowski K, Tomchick DR, et al. Bacterial pseudokinase catalyzes protein polyglutamylation to inhibit the SidE-family ubiquitin ligases. Science. 2019;364: 787–792. doi:10.1126/science.aaw7446

36. Ingmundson A, Delprato A, Lambright DG, Roy CR. Legionella pneumophila proteins that regulate Rab1 membrane cycling. Nature. 2007;450: 365–9. doi:10.1038/nature06336

37. Ham H, Orth K. De-AMPylation unmasked: modulation of host membrane trafficking. Sci Signal. 2011;4: pe42. doi:10.1126/scisignal.2002458

38. Iyer S, Das C. The unity of opposites: Strategic interplay between bacterial effectors to regulate cellular homeostasis. J Biol Chem. 2021;297: 101340. doi:10.1016/j.jbc.2021.101340

39. Elez K, Bonvin AMJJ, Vangone A. Distinguishing crystallographic from biological interfaces in protein complexes: role of intermolecular contacts and energetics for classification. BMC Bioinformatics. 2018;19: 438. doi:10.1186/s12859-018-2414-9

40. Fu J, Li S, Guan H, Li C, Zhao Y-B, Chen T-T, et al. Legionella maintains host cell ubiquitin homeostasis by effectors with unique catalytic mechanisms. Nat Commun. 2024;15: 5953. doi:10.1038/s41467-024-50311-2

41. Gross SR, Kinzy TG. Improper organization of the actin cytoskeleton affects protein synthesis at initiation. Mol Cell Biol. 2007;27: 1974–89. doi:10.1128/MCB.00832-06

42. Kim S, Coulombe PA. Emerging role for the cytoskeleton as an organizer and regulator of translation. Nat Rev Mol Cell Biol. 2010;11: 75–81. doi:10.1038/nrm2818

43. Lenk R, Ransom L, Kaufmann Y, Penman S. A cytoskeletal structure with associated polyribosomes obtained from HeLa cells. Cell. 1977;10: 67–78. doi:10.1016/0092-8674(77)90141-6

44. Kaminska M, Havrylenko S, Decottignies P, Le Maréchal P, Negrutskii B, Mirande M. Dynamic Organization of Aminoacyl-tRNA Synthetase Complexes in the Cytoplasm of Human Cells. J Biol Chem. 2009;284: 13746–13754. doi:10.1074/jbc.M900480200

45. Stapulionis R, Kolli S, Deutscher MP. Efficient mammalian protein synthesis requires an intact F-actin system. J Biol Chem. 1997;272: 24980–6. doi:10.1074/jbc.272.40.24980

46. Fontana MF, Banga S, Barry KC, Shen X, Tan Y, Luo Z-Q, et al. Secreted bacterial effectors that inhibit host protein synthesis are critical for induction of the innate immune response to virulent Legionella pneumophila. PLoS Pathog. 2011;7: e1001289. doi:10.1371/journal.ppat.1001289

47. Belyi Y. Targeting Eukaryotic mRNA Translation by Legionella pneumophila. Front Mol Biosci. 2020;7: 80. doi:10.3389/fmolb.2020.00080

48. De Leon JA, Qiu J, Nicolai CJ, Counihan JL, Barry KC, Xu L, et al. Positive and Negative Regulation of the Master Metabolic Regulator mTORC1 by Two Families of Legionella pneumophila Effectors. Cell Rep. 2017;21: 2031–2038. doi:10.1016/j.celrep.2017.10.088

49. Chauhan D, Joseph AM, Shames SR. Intrabacterial Regulation of a Cytotoxic Effector by Its Cognate Metaeffector Promotes Legionella pneumophila Virulence. mSphere. 2023; e0055222. doi:10.1128/msphere.00552-22

50. Wang T, Song X, Tan J, Xian W, Zhou X, Yu M, et al. Legionella effector LnaB is a phosphoryl-AMPylase that impairs phosphosignalling. Nature. 2024;631: 393–401. doi:10.1038/s41586-024-07573-z

51. Chen T-T, Lu Q, Zheng S-R, Fu J, Chen J, Kang L, et al. Structure and mechanism of an actin-dependent bacterial phosphoryl AMPylase. Nat Chem Biol. 2025. doi:10.1038/s41589-025-01945-w

